# Path of differentiation defines human macrophage identity

**DOI:** 10.1101/2025.01.24.634694

**Authors:** Katie Frenis, Brianna Badalamenti, Ohanez Mamigonian, Chen Weng, Dahai Wang, Sara Fierstein, Parker Côté, Hesper Khong, Hojun Li, Edroaldo Lummertz da Rocha, Vijay G. Sankaran, R. Grant Rowe

**Author notes:** **CORRESPONDENCE:** Grant Rowe, 1 Blackfan Circle, Karp 7, Boston MA 02115.

## Abstract

Macrophages play central roles in immunity, wound healing, and homeostasis - a functional diversity that is underpinned by varying developmental origins. The impact of ontogeny on properties of human macrophages is inadequately understood. We demonstrate that definitive human fetal liver (HFL) hematopoietic stem cells (HSCs) possess two divergent paths of macrophage specification that lead to distinct identities. The monocyte-dependent pathway exists in both prenatal and postnatal hematopoiesis and generates macrophages with adult-like responses properties. We now uncover a fetal-specific pathway of expedited differentiation that generates tissue resident-like macrophages (TRMs) that retain HSC-like self-renewal programs governed by the aryl hydrocarbon receptor (AHR). We show that AHR antagonism promotes TRM expansion and mitigates inflammation in models of atopic dermatitis (AD). Overall, we directly connect path of differentiation with functional properties of macrophages and identify an approach to promote selective expansion of TRMs with direct relevance to inflammation and diseases of macrophage dysfunction.

## INTRODUCTION

Macrophages fulfill critical roles in organogenesis, immunity, macromolecule metabolism, and wound healing during both development and homeostasis. They are exquisitely sensitive to microenvironmental cues, interpreting signals from their tissue of residence to shape their identity and behavior^1–3^. Another layer of macrophage diversity lies in their origins; while it was historically believed that TRMs are continuously replenished by monocytes produced from HSCs, more recent studies using murine lineage tracing models have demonstrated that many TRM subsets arise from pre-definitive hematopoietic waves and stably populate tissues via self-renewal^4–6^. In contrast to this model, human HSC transplantation has demonstrated that TRMs can be facultatively replaced from allografted definitive adult HSCs, indicating that human macrophage ontogeny may be plastic and/or depart from paradigms established in mice^7,8^.

The self-renewing potential of embryonically-derived TRMs stands in contrast to most other terminally differentiated hematopoietic cells, including monocytes and monocyte-derived macrophages (moMacs)^9,10^. In normal hematopoiesis, self-renewal is generally restricted to HSCs and proliferative capacity wanes during differentiation. How TRMs retain self-renewal potential is poorly understood; possibilities include engagement of a lineage-specific programs as occurs in lymphocytes or appropriation of HSC-like programs preserved or reactivated through differentiation^11^. In the latter case, at least some TRMs would be hypothesized to be of definitive origin due to the lack of long-term self-renewal in primitive hematopoietic progenitors^12^. Though stimulation by macrophage colony stimulating factor (M-CSF), a critical cytokine for TRMs, is linked to long-term survival of macrophages, little is known about how self-renewal programs are regulated in human TRM specification^13,14^.

TRMs form interstitial networks in every organ, positioning them for rapid response to tissue compromise^15^. They initiate acute inflammation by recruiting effector inflammatory cells, including moMacs, from the blood. Notably, fetal-derived TRMs are absent in acute inflammation after their initial response, only repopulating during resolution^15^. Though they are positioned to carry out inflammation, TRMs yield to moMacs as the driving force. This suggests that macrophage heterogeneity could, in part, arise from the identity of the originating cell. Further supporting this, cord blood and neonatal monocytes exhibit relative insensitivity to TLR4 activation and do not assume adult-like inflammatory responses until the first year of life^27,28^. This homeostatic TRM:moMac balance is disrupted in many diseases such as AD, atherosclerosis, graft-versus-host disease, and arthritis^18–25^. While it is accepted that macrophages adjust their polarization state in response to their environment and that moMacs can assume a TRM-like identity in certain contexts, macrophage heterogeneity within the microenvironment suggests that a factor additional to outside of environment can shape macrophage phenotype^2,14,24–26^.

Here, predicated upon gene expression profiling suggesting that definitive HFL HSCs possess TRM potential, we examined macrophage specification in HFL definitive HSCs relative to postnatal HSCs. We found that while both types of HSCs could generate moMacs, only HFL HSCs possessed an alternative expedited pathway of TRM specification. This occurred due to apparent priming of macrophage fate in a subset of HFL HSCs, enabling rapid acquisition of TRM identity by bypassing a monocyte intermediate. This relatively direct change in cell state results in retention of elements of AHR-regulated HSC self-renewal programs, enabling selective expansion of anti-inflammatory TRMs in models of AD. Overall, our findings elucidate a developmentally restricted pathway of human TRM specification from definitive HSCs that can potentially be therapeutically regulated.

## RESULTS

### Expedited differentiation of fetal multipotent HSPCs to macrophages

In a published atlas of human HSPCs across the lifespan, we observed that adult HSPCs expressed monocyte-associated transcripts *S100A4*, *S100A6*, *S100A8*, *S100A9*, *ANXA1*, and *LYZ,* as expected of monocytic potential^29^. In contrast, fetal HSPCs expressed transcripts associated with TRM identity: *C1QA, HMOX1, MAF*, *LIPA*, *CD163*, *MAFB*, *LYVE1*, and *FOLR2* **(Figure 1A)**^29–31^. These contrasting profiles supported TRM potential in human HSPCs at early gestational ages (10-15 PCW) and led us to hypothesize that macrophage differentiation was age-dependently programmed at the HSPC level^31^. We speculated that expression in HSPCs at this temporal stage could indicate a differentiation program specific to HFL hematopoiesis^32–34^.

**Figure 1:**
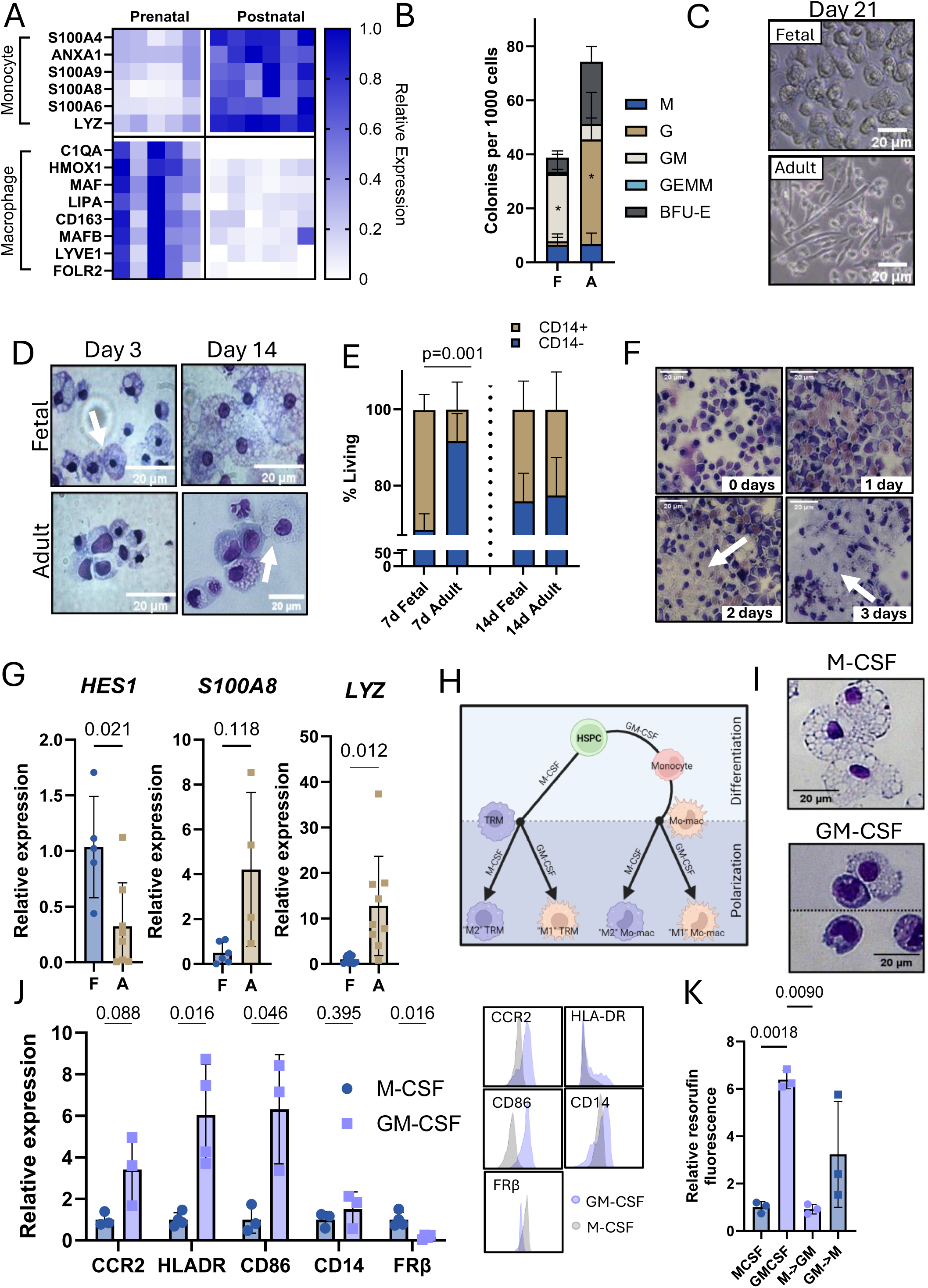
Expedited differentiation of fetal multipotent HSPCs to macrophages. **A.** Heatmap of selected monocyte and macrophage genes expressed at the HSPC level, curated from Mulder et al^29^. Data from Li et al^31^. Each column represents one donor: n = 5 human fetal liver donors (10-15 PCW); n = 6 adult bone marrow donors (25-53 years old)^31^. **B.** Proportions of colony-forming units (CFUs) scored in CFU assays. Data are presented as mean ± SD. n = 4 biological replicates per age; two-way ANOVA with Bonferroni’s correction. * indicates p < 0.05. **C.** Phase contrast image of mature differentiated macrophages in culture. Magnification: 200x; scale bars = 20 µm. **D.** Cytospins of fetal and adult cultures after 3 and 14 days in M-CSF-driven differentiations. Arrows indicate the first appearance of mature macrophages. Magnification: 200x; scale bars = 20 µm. **E.** In vitro macrophage commitment after 7 and 14 days in culture. Data are presented as mean ± SD. Tan bars indicate proportion of living cells expressing CD14. n = 4 biological replicates per age; two-way ANOVA with Bonferroni’s correction. **F.** Images of daily sampling of fetal differentiation. Labels indicate number of days in culture with M-CSF; arrows highlight macrophages. Magnification: 100x; scale bars = 20 µm. **G.** Expression of HES1, S100A8, and LYZ by qPCR. Bars represent fold change calculated from average fetal expression ± SD. n = 5-9 biological donors per age; two-tailed, unpaired t-tests with Welch’s correction. **H.** Differentiation scheme to produce fetal monocyte-derived macrophages. After differentiation with M-CSF or GM-CSF, mature macrophages were repolarized as indicated to equilibrate cell state. **I.** Representative images of cells after 14 days of differentiation with monocytes present in the GM-CSF differentiation. Magnification: 200x; scale bars = 20 µm. **J.** Surface marker analysis via flow cytometry of HFL M-CSF versus GM-CSF differentiated macrophages. Bars represent fold change calculated from average fetal expression ± SD. n = 3-4 donors; unpaired two-tailed t-tests. **K.** Peroxidase activity of fMac^M-CSF^or fMac^GM-CSF^ macrophages. M-CSF/GM-CSF indicates the macrophages were differentiated and maintained in culture with the respective cytokine. M->GM and GM->M indicate the driver of differentiation (first) and the cytokine used to repolarize (second). Represented as the relative fluorescent intensity ± SD. n = 3 donors, multiple paired t-tests.

Based on this analysis, we examined macrophage potential in HSCs and multipotent progenitors (MPPs) sorted from HFL and adult bone marrow (ABM) (Lineage^−^CD34^+^CD38^−^, **Supplemental Figure 1A-B**) in colony forming unit (CFU) assays. Although HFL HSPCs produced relatively more bipotent granulocyte-monocyte colonies than adult HSPCs, there was not an increase in unilineage monocytic colonies **(Figure 1B)**. We then hypothesized that macrophage priming could be growth factor dependent, as only granulocyte-macrophage colony stimulating factor (GM-CSF), but not M-CSF is included in standard CFU medium. Since GM-CSF produces inflammatory “M1” macrophages while M-CSF produces “M2” macrophages, a common state for TRMs, we performed M-CSF-driven differentiation in liquid culture^33,35^. HFL macrophages were amoeboid, and lightly adherent with many proliferative clusters, while ABM macrophages cultures exhibited classical features: strong adherence and spindle-shaped morphology **(Figure 1C, Supplemental Figure 1C)**. HFL macrophages were evident within the first week of culture, whereas ABM macrophages required 14 days to reach equivalent CD14^+^ proportions, indicating stage-specific kinetics of differentiation **(Figure 1D-E).** In daily sampling of fetal differentiation, a monocyte stage was not detected before the first macrophages emerged **(Figure 1F, Supplemental Figure 1D).** We measured the expression of monocyte-associated genes in terminal macrophages and observed that HFL macrophages had high expression of *HES1,* a marker of self-renewing TRMs, but low expression of monocyte genes *LYZ* and *S100A8* relative to adult **(Figure 1G)**^29^. To enable comparison between rapidly differentiating macrophages and those arising via a monocyte intermediate, we split fetal HSPCs to differentiate with either M-CSF (fMac^M-CSF)^ or GM-CSF (fMac^GM-CSF^)and after 14 days, selected terminal macrophages to be cultured for five additional days with either M-CSF (fMac^M->GM^) or GM-CSF (fMac^GM->M^) to equilibrate polarization state **(Figure 1H)**. fMac^GM-CSF^ differentiated through a monocyte intermediate, confirming that HFL HSPCs could produce macrophages via the canonical path **(Figure 1I)**. fMac^GM-CSF^ exhibited an activated moMac immunophenotype with higher expression of CCR2, HLA-DR, and CD86 than fMac^M-CSF^, which was defined by high expression of folate receptor β (FRβ), a marker of human self-renewing macrophages **(Figure 1J)**^36^. fMac^GM-CSF^ were repolarized with M-CSF (fMac^GM->M^) and expectedly downregulated CD86 but maintained higher HLA-DR and lower FRβ expression as well as higher peroxidase activity than fMac^M-CSF^ and its counterpart fMac^GM->M^, indicating a lasting effect from path of differentiation **(Supplemental Figure 1E, Figure 1K)**^36–39^. These findings indicate that fetal HSPCs can generate macrophages through a monocyte-independent pathway under M-CSF-driven differentiation while retaining access to the canonical monocyte-dependent differentiation. Because fMac^M-CSF^ resembled TRMs and apparently bypassed the monocyte state as evidenced by rapid differentiation and lack of acquisition of monocyte markers, we then hypothesized that expedited, monocyte-independent differentiation specifies TRMs in definitive hematopoiesis^29^.

### Fetal HSPCs produce tissue resident-like macrophages in culture and in vivo

In M-CSF driven assays, ABM macrophages (aMac^M-CSF^) resembled fMac^GM-CSF^ with low expression of FRβ and higher expression of CCR2, HLA-DR, and CD86, further suggesting that transit through a monocyte intermediate permanently impacted the macrophage’s downstream phenotype **(Figure 2A)**. In native HFL and ABM tissue, we detected both FRβ^high^ and FRβ^low^ macrophages, though FRβ^high^ macrophages were enriched in the HFL **(Figure 2B, Supplemental Figure 2A).** In both fractions, HLA-DR was higher in ABM macrophages than HFL macrophages, while the reverse was observed for FRβ, validating our immunophenotyping in cultured macrophages (**Figure 2C-D**). In line with our observations of fMac^M-CSF^ possessing a self-renewing macrophage immunophenotype, these macrophages cycled more than adult counterparts and maintained downregulated *MAF* and *MAFB* with upregulated *MYC*, a state reported to enable terminal macrophages to self-renew **(Figure 2E-F)**^40^. To validate the observed stage-specific mechanisms of macrophage differentiation, we xenotransplanted HFL or ABM HSPCs into NOD.Cg-Prkdc^scid^Il2rg^tm1Wjl^Tg(CMV-IL3,CSF2,KITLG)1EavTg(CSF1)3Sz/J (NSG-Q) mice expressing transgenic human stem cell factor (SCF), interleukin-3 (IL-3), M-CSF and GM-CSF^41^. After four weeks, we observed higher overall multilineage human chimerism from HFL HSPCs **(Supplemental Figure 2B-F)**. While adult HSPCs produced higher relative proportion of monocytes in the xenotransplanted bone marrow, HFL HSPCs readily produced FRβ^+^ TRMs in the liver and spleen, tissues where macrophages are canonically fetal-derived **(Figure 2G-I, Supplemental Figure 2G)**^6,42^. However, we did not observe significant differences in FRβ^+^TRM presence in the peritoneum and heart, where TRMs are partially monocyte-derived, suggesting that high fetal chimerism in the liver and spleen was not simply a consequence of higher engraftment **(Figure 2H-I)**^43–45^. Overall, upon xenotransplantation, fetal HSPCs exhibited relatively low baseline monocyte potential but high potential for producing TRMs, which also exhibited low peroxidase activity **(Supplemental Figure 2G-H)**. Together, these observations informed our hypothesis that fetal macrophages have access to a second monocyte-independent differentiation pathway that specifies phenotypically distinct TRMs.

**Figure 2:**
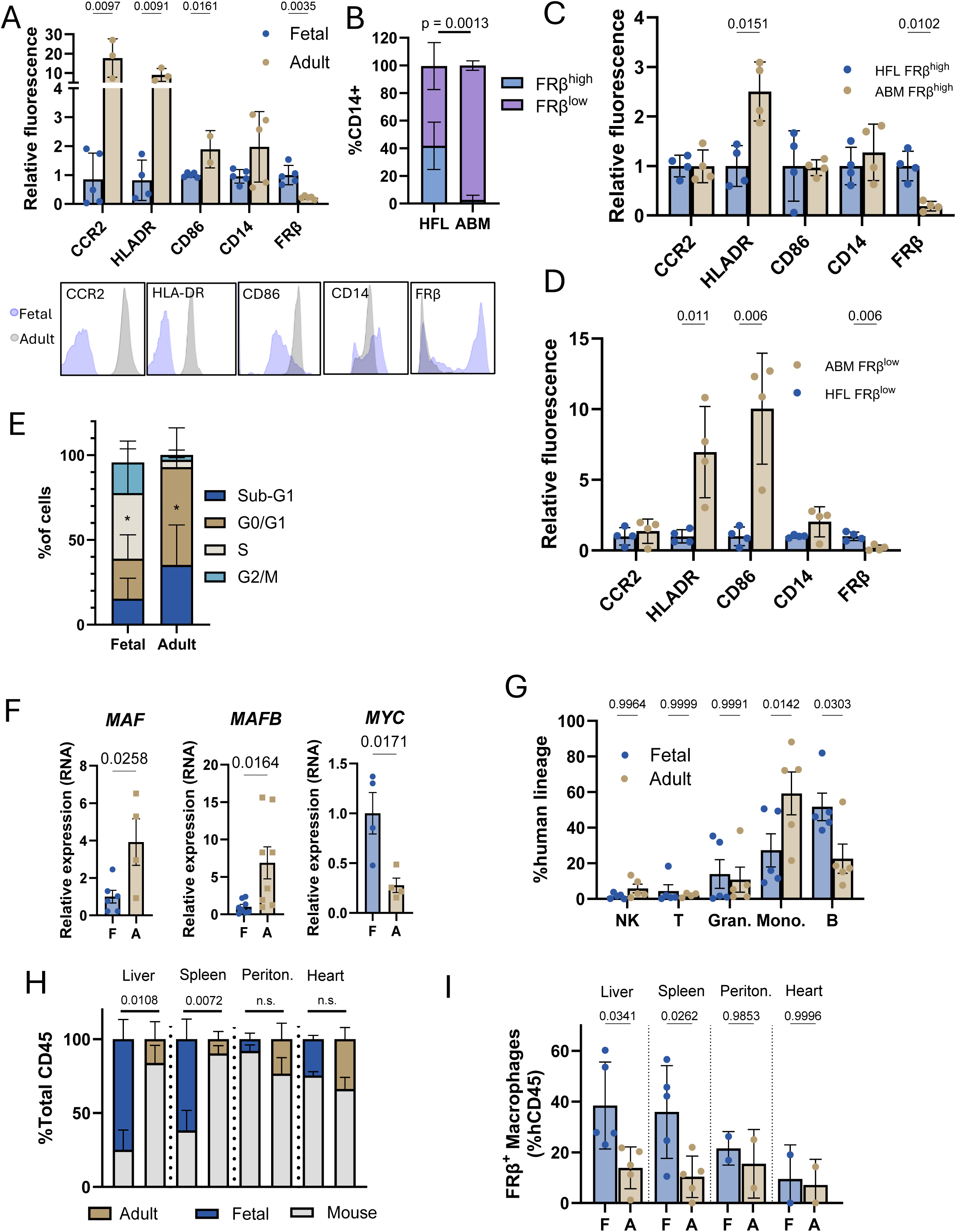
Fetal HSPCs produce tissue resident-like macrophages in culture and in vivo. **A.** Immunophenotype of cultured HFL and ABM macrophages after approximately 21 days in culture, measured by flow cytometry. Mean fluorescent signal (MFI) was normalized to the average of the fetal signal. Representative histograms are shown below. n = 2-5 donors per age; multiple unpaired t-tests with Holm-Sidak correction and an alpha of 0.05. **B.** Proportions of FRβ^high^and FRβ^low^ macrophages in native HFL and ABM. n = 4 donors per age; 2-way ANOVA with Sidak’s multiple comparisons test. **C-D.** Immunophenotype of FRβ^high^and FRβ^low^ macrophages in HFL and ABM, respectively. MFI was measured by flow cytometry and normalized to the average of the fetal signal to represent fold difference ± SD. n = 4 donors per age; multiple unpaired t-tests with Holm-Sidak correction and an alpha of 0.05. **E.** Cell cycle analysis of fetal and adult M-CSF cultured macrophages, measured by flow cytometric measurement of Ki67 with propidium iodide. n = 5 fetal; n = 4 adult. Error bars represent SD. Representative plots are shown to the right. * indicates p < 0.05. **F.** Expression of MAF, MAFB, and MYC by qPCR. Bars represent fold change calculated from average fetal expression ± SD. n = 4-8 biological donors per age; two-tailed, unpaired t-tests. **G.** Lineage composition in the bone marrow of NSG-Q mice. Bars represent the percent of the human lineage compartment (the sum of cells expressing human CD45 and a lineage marker, CD3, CD14, CD19, CD16, CD56) ± SD. NK; natural killer, T; T cells, Gran; granulocytes, Mono; monocytes, B; B cells. n = 5 donors per age; two-way ANOVA with Sidak’s multiple comparisons test. **H.** Human chimerism in liver, spleen, heart, and peritoneum as percent of total CD45 ± SD. n = 5 donors per age for liver and spleen; n = 2 donors per age for peritoneum and heart. Multiple unpaired t-tests with Holm-Sidak correction and an alpha of 0.05. **I.** FRβ^+^ macrophages as a percent of human CD45^+^ cells ± SD. F indicates fetal transplanted cells, A indicates adult. n = 5 biological replicates per age for liver and spleen; n = 2 biological replicates per age for peritoneum and heart. One-way ANOVA with Sidak’s multiple comparisons test.

### HFL HSCs possess two trajectories of macrophage specification

To directly investigate mechanisms of developmental differences in human macrophage specification, we performed single cell RNA sequencing (scRNAseq) on native human macrophages and defined upstream progenitors isolated from native hematopoietic tissues **(Supplemental Figure 3A-C)**. Following Louvain clustering, we annotated each cluster using hallmark genes along differentiation pathways and generated cluster signatures via differential gene expression **(Figure 3A-C; Supplemental Figure 4A-B, Supplemental Tables 1-2)**. We identified multiple distinct transcriptional clusters of macrophages and intermediates at both ages as well as contaminating lymphocytes (Lymph1^ABM^/Lymph2^ABM^) in ABM. In HFL, there were two readily identifiable macrophage populations: one resembling self-renewing TRMs that clustered independently (TRM^HFL^) and another that clustered adjacent to monocytes (moMac^HFL^) in UMAP space (**Figure 3A)**. There were three apparent macrophage populations in the ABM that we identified by gene expression: moMac^ABM^, erythroblast island macrophages (EBI^ABM^), and MHC-II-expressing macrophages (MHC-II^ABM^; **Figure 3B; Supplemental Figure 4B)**^46,47^. Compared to all macrophage populations, TRM^HFL^ and moMac^HFL^ showed the highest expression of “M2”-associated transcripts *CD200R1* and *CD163*, respectively (**Figure 3D**). All ABM macrophages and moMac^HFL^ had expected core moMac signatures of *PTPRC, ITGAM*, *CD38*, *CD44*, *CD74*, and *ITGAX* (**Figure 3D**). Though TRM^HFL^ had low expression of these markers, its expression of *CD14* was the highest and expression of self-renewal related TFs *FLI1*, *MECOM*, *ERG*, *ID1*, and *MYC* was uniquely high **(Figure 3D)**. Application of signatures of defined macrophage subsets revealed that TRM^HFL^ resembled the *HES1* cluster from Mulder et al, which is notably described as self-renewing TRMs with dissimilarity to monocytes^29^, while moMac^HFL^ resembled the C1Q^high^ population, an “M2” cluster that is likely monocyte-derived **(Figure 3E)**. ABM macrophages all appeared monocyte-derived based on their expression of monocyte signature genes **(Figure 3E)**^29^.

**Figure 3:**
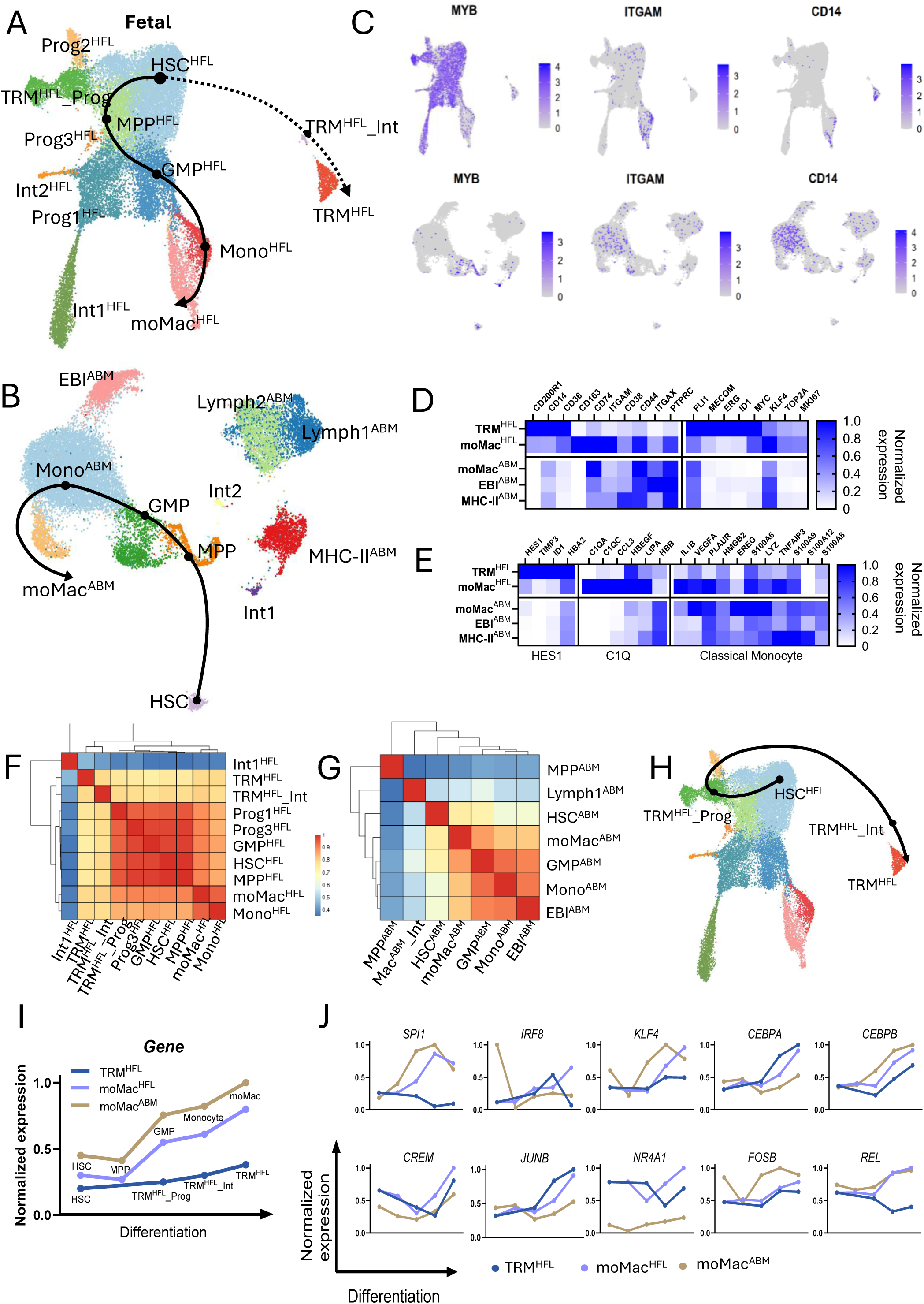
Fetal HSCs possess two modes of macrophage specification. **A-B.** UMAP projection of fetal and adult cells, respectively. All clusters are annotated on the UMAP; canonical pathways are marked with black lines. The hypothesized fetal-specific pathway is marked with a dotted line. Points indicate discrete populations along differentiation pathways. **C.** Highlight plots of key marker genes (MYB, ITGAM, CD14) used to identify clusters, fetal on top and adult on the bottom. **D.** Expression comparison of all identified macrophage clusters. Benchmark surface markers on the left of the heatmap illustrate differences in key macrophage-associated genes. On the right, HSC-associated self-renewal genes are shown. **E.** Curated genes used to identify macrophage identities, annotated with the cluster names from Mulder et al below^29^. On the left, genes associated with fetal-derived self-renewing macrophages (HES1 macrophages), and fetal moMacs (C1Q macrophages) are shown. On the right, curated genes from profiles of classical monocytes are shown^29^. **F-G.** Similarity indices generated from the top 2000 genes that exhibited the highest variance within all clusters with the indicated comparisons. **H.** Differentiation pathway generated by CellRouter when HSCs were input as the initiating cluster and TRMs as the endpoint^48^. **I.** Model for interpretation of gene expression analysis across differentiation. The pseudobulk value for each cluster was normalized to a maximum of 1 (y-axis) and plotted to include populations along the canonical and projected differentiation pathways (x-axis) to trace expression changes over the course of differentiation. Canonical differentiations generating moMac^HFL^ and moMac^ABM^ included 5 discrete clusters (from left to right: HSC – MPP – GMP – Monocyte – Macrophage), whereas 4 were included in TRM^HFL^ differentiation (from left to right: HSC – TRM^HFL^_Prog – TRM^HFL^_Int – TRM). **J.** Expression of selected TFs along the indicated trajectories. Blue; TRM^HFL^, lavender. moMac^HFL^, tan; moMac^ABM^.

Because moMac^ABM^ bears strong similarity to moMac^HFL^, is reportedly present in many human tissues, and was not a ‘specialized’ bone marrow macrophage, we used this cluster as the adult moMac benchmark population for further analyses^36^. Using the top 2000 most variable genes across the clusters, we generated similarity indices amongst clusters, revealing expectedly high similarity between Mono^HFL^ and moMac^HFL^ as well as Mono^ABM^ and moMac^ABM^ (**Figure 3F-G**). TRM^HFL^, however, only had a moderate transcriptional relationship with Mono^HFL^ and moMac^HFL^ **(Figure 3F-G)**. We next performed trajectory analysis to define a differentiation trajectory from HSC to TRM^HFL^ with the intention of identifying TFs that contribute to the specification of the self-renewing TRM population^48^. CellRouter identified a pathway in HFL without the conventional myeloid progenitor and monocyte intermediates, but rather included a *CD34*^+^*CD38*^+^*MPO*^−^ cluster (TRM^HFL^_Prog) that did not resemble classic granulocyte-monocyte progenitors (GMPs) and a *CD38*^+^*MYC*^+^*CD34*^low^*CD14*^low^*MPO*^−^ intermediate (TRM^HFL^_Int; **Figure 3H, Supplemental Figure 4A)**.

To compare this differentiation pathway with the canonical monocyte-dependent pathway, we curated a list of TFs known to coordinate monocyte/macrophage differentiation and plotted their expression along these state change trajectories. Expression of *SPI1*, *IRF8*, and *KLF4*, the master regulators of monocyte differentiation, was acquired as expected in moMac^HFL^ and moMac^ABM^ differentiation, in contrast to the TRM^HFL^ path where these TFs were not engaged **(Figure 3I-J)**.TRM^HFL^ also did not acquire expression of *REL*, a TF in monocyte development with an important role in inflammatory action **(Figure 3J)**^49–51^. Conversely, TRM^HFL^ gained expression of TFs that coordinate anti-inflammatory and homeostatic status: *CREM*, *JUNB*, *NR4A1*, and *CEBPA* (**Figure 3J**)^49,52,53^. Together, these findings support a monocyte-independent, fetal-specific pathway of TRM specification in native human hematopoiesis and suggest this is an expedited means toward production of TRMs in a narrow developmental time window.

### A developmentally-restricted, macrophage primed HFL HSC

Though some lineage tracing studies in mice have posited a primitive hematopoietic origin of TRMs, other murine studies have suggested definitive origins and definitive fetal HSCs have been reported to express markers of macrophage potential^1,54,55^. To determine whether such mechanisms underlie HFL-specific, HSC-level TRM priming, we examined expression of receptors for growth factors on HSCs from the current dataset and our dataset of HSPCs across the lifespan **(Supplemental Figure 5A-B)**^31^. We observed a subset of HFL, but not ABM, HSCs and MPPs expressing *CSF1R*, encoding the receptor for M-CSF **(Figure 4A, Supplemental Figure 5C)**. Using flow cytometry, we also detected a subset of immunophenotypic definitive HFL HSCs (defined as Lineage^−^CD34^+^CD38^−^CD90^+^) with surface expression of CSF1R that was much more abundant than ABM counterparts **(Figure 4B-C)**. To evaluate whether this immunophenotype conferred TRM priming, we sorted single CSF1R^+^ or CSF1R^−^ HFL or ABM HSCs for M-CSF-driven differentiation. Starting on day two after sorting, the wells were scored for viability and differentiation. We observed that HFL CSF1R^+^ HSCs had a viability advantage in the presence of M-CSF **(Figure 4D).** Approximately 60% of the surviving HFL CSF1R^+^ HSCs showed evidence of macrophage differentiation within five days **(Figure 4E).** To confirm definitive hematopoietic potential (i.e., myeloid/lymphoid multipotency), we then sorted single CSF1R^+^ and CSF1R^−^ clones into MS5 stromal culture with interleukin 7 to drive B-lymphocyte development with no observed differences in lymphoid potential after 21 days **(Figure 4F)**. We then xenotransplanted CSF1R^+^ or CSF1R^−^ cells into immunodeficient mice and observed lymphoid differentiation from both sources and fetal-type hemoglobin **(Figure 4G-H**). In our scRNAseq dataset, the fetal HSC cluster expressed definitive *HOX* genes, and expression of *MECOM*, *PROCR*, and *HLF*, and other genes that have been shown to be expressed in emerging HSCs (**Supplemental Figure 5D-E**)^56–60^. Overall, we interpreted these profiles to indicate that these TRM-primed multipotent HSCs are definitive in origin **(Figure 4H; Supplemental Figure 5D-E)**.

**Figure 4:**
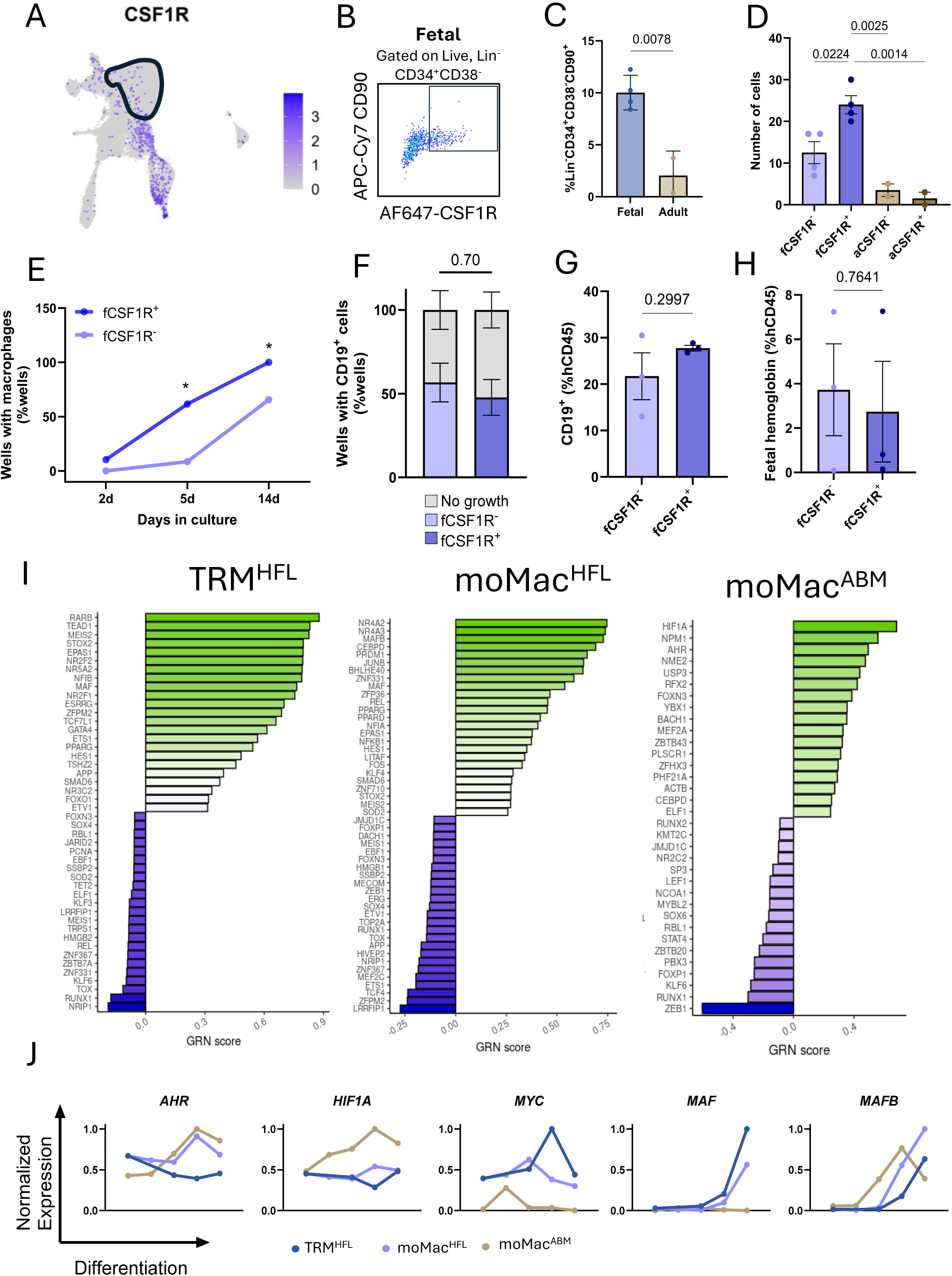
Priming of TRM fate in definitive HFL HSCs. **A.** Expression of CSF1R within the fetal UMAP. The fetal HSC cluster is circled; blue indicates cells expressing CSF1R. **B.** Representative plot of fetal Lin^−^CD34^+^CD38^−^CD90^+^ HSCs expressing CSF1R, measured by flow cytometry. **C.** Quantification of abundance of CSF1R^+^ HSCs as a fraction of the HSC compartment in both HFL and ABM ± SD. n = 4 for HFL and n = 1 for ABM. Unpaired two-tailed t-test. Points represent biological replicates. **D.** Day 2 post-FACS viability. Single cells were sorted into wells in the presence of M-CSF and manually scored on the second day for live outgrowths. Bars represent percentage of wells with living cells ± SD. n = 4 for fetal, n = 2 for adult; One-way ANOVA with Tukey’s multiple comparison’s test. **E.** Following single cell sorts of CSF1R^−^ HSC and CSF1R^+^ HSC, wells were blindly scored for differentiation status at 2, 5, and 14 days. Bars represent the percent of wells with a differentiated macrophage ± SD. n = 4 biological HFL replicates; two-way ANOVA with Bonferroni’s correction. * indicates p < 0.05. **F.** single CSF1R+ and CSF1R-clones were sorted into culture with IL-7 to drive lymphoid development. CSF1R^+^ and CSF1R^−^ clones showed no difference in lymphoid output or survival. Data is presented as the percent of sorted wells. Gray indicates no outgrowth, lavender and blue indicate detection of CD19^+^ cells after 21 days. **G-H.** 2000 CSF1R^+^ and CSF1R^−^ HSCs were transplanted directly into the femur marrow of NSG mice. Bars represent CD19 expression and fetal hemoglobin as a percent of hCD45 ± SD. n = 3 fetal donors; two-tailed, unpaired t-tests. **I.** TF behavior over TRM^HFL^, moMac^HFL^, and moMac^ABM^ differentiation. Length of the bar corresponds to change in expression over the course of differentiation. **J.** Scaled expression of the indicated TFs over each differentiation pathway. Clusters in the trajectory are plotted on the x-axis from left to right (points), least to most differentiated. Pseudobulk expression value is plotted on the y-axis.

M-CSF signaling interfaces with several pathways that regulate the expression of many TFs, so we returned to the scRNAseq dataset to narrow M-CSF targets that could be driving differentiation. We analyzed TF behavior over the course of canonical and expedited differentiations and generated lists of TFs that were up- and down-regulated in each macrophage differentiation **(Figure 4I)**. We also conducted enrichment analysis by Enrichr and GSEA of these TFs and their regulons **(Supplemental Table 3)**. Expression of *MAF*, encoding a TF induced by M-CSF, was shared in the upregulated TFs of TRM^HFL^ and moMac^HFL^ pathways, but *MAFB* was only observed in moMac^HFL^ (**Figure 4I**). *AHR*, *HIF1A*, *FOXO3*, *FOXO1*, *FOS*, and *BACH1* were upregulated in the moMac^ABM^ pathway (**Figure 4I)**. Several of the identified TFs have defined roles in self-renewal; *AHR* negatively regulates self-renewal in HSPCs, downregulates *MYC*, and appears to be a hallmark of monocyte identity^29,61^. *MAFB* restrains self-renewal and induces monocyte/macrophage differentiation^40,62–64^. Finally, *HIF1A* participates in a feedback loop with *MYC,* an important driver of self-renewal in macrophages, and coordinates inflammatory programs, which are known to inhibit self-renewal^50,51,65–67^. We then plotted the scaled expression of these TFs over the course of the three differentiation pathways **(Figure 4J)**. TRM^HFL^ did not acquire *HIF1A* or *AHR* over the course of differentiation but moMac^HFL^ and moMac^ABM^ had highest expression at the monocyte state, which was then retained in the subsequent macrophage (**Figure 4J**). *MYC*, a known driver of macrophage proliferation, steeply reduced preceding the monocyte stage, suggesting that transit through the monocyte intermediate restricts moMac^HFL^ and moMac^ABM^ self-renewal, which is retained by TRM^HFL^ via expedited differentiation **(Figure 4J)**.

### Fetal TRMs are homeostatic and anti-inflammatory relative to moMacs

In general, TRMs perform homeostatic and trophic tasks rather than execute acute inflammation, which is a responsibility of moMacs^68–70^. moMac^HFL^ and moMac^ABM^ shared high expression of Fc receptors, HLA genes, costimulatory molecules, and TLR/NLR receptors, indicative of preparation to respond to pathogens or tissue damage, as would be expected of a moMac (**Figure 5A**). While TRM^HFL^ has low expression of these markers, it expressed higher levels of scavenger receptors *CD36* and *COLEC12*, as would be expected of a homeostatic TRM primed for metabolism and clearance of lipoproteins **(Figure 5A).** A comparison of the 5000 highest expressed genes showed that TRM^HFL^ possessed a distinct profile, whereas many genes were shared by monocytes, moMac^HFL^ and moMac^ABM^ **(Figure 5B)**. When quantified as percentages, TRM^HFL^ only shared 68.14% and 73.78% of the top 5000 genes with moMac^ABM^ and moMac^HFL^, respectively. Of those genes shared, 56.46% were shared by all clusters **(Figure 5C)**. To better understand how these macrophages genetically diverged, we performed KEGG pathway analysis of the genes that were solely expressed in each macrophage cluster and further confirmed that many of the characteristic pathways of moMac^HFL^ and moMac^ABM^ were immune-related, whereas TRM^HFL^ signature was more homeostatic in nature **(Figure 5D)**. To confirm these findings, we conducted functional phagocytosis assays. aMac^M-CSF^ had comparatively higher phagocytic capacity than fMac^M-CSF^, despite identical differentiation and culture conditions **(Figure 5E)**. fMac^GM-CSF^, which differentiated through a monocyte intermediate, exhibited higher phagocytic response and reactive oxygen species (ROS) production compared to donor-matched fMac^M-CSF^, regardless of induced polarization state (**Figure 5F-G)**.

**Figure 5:**
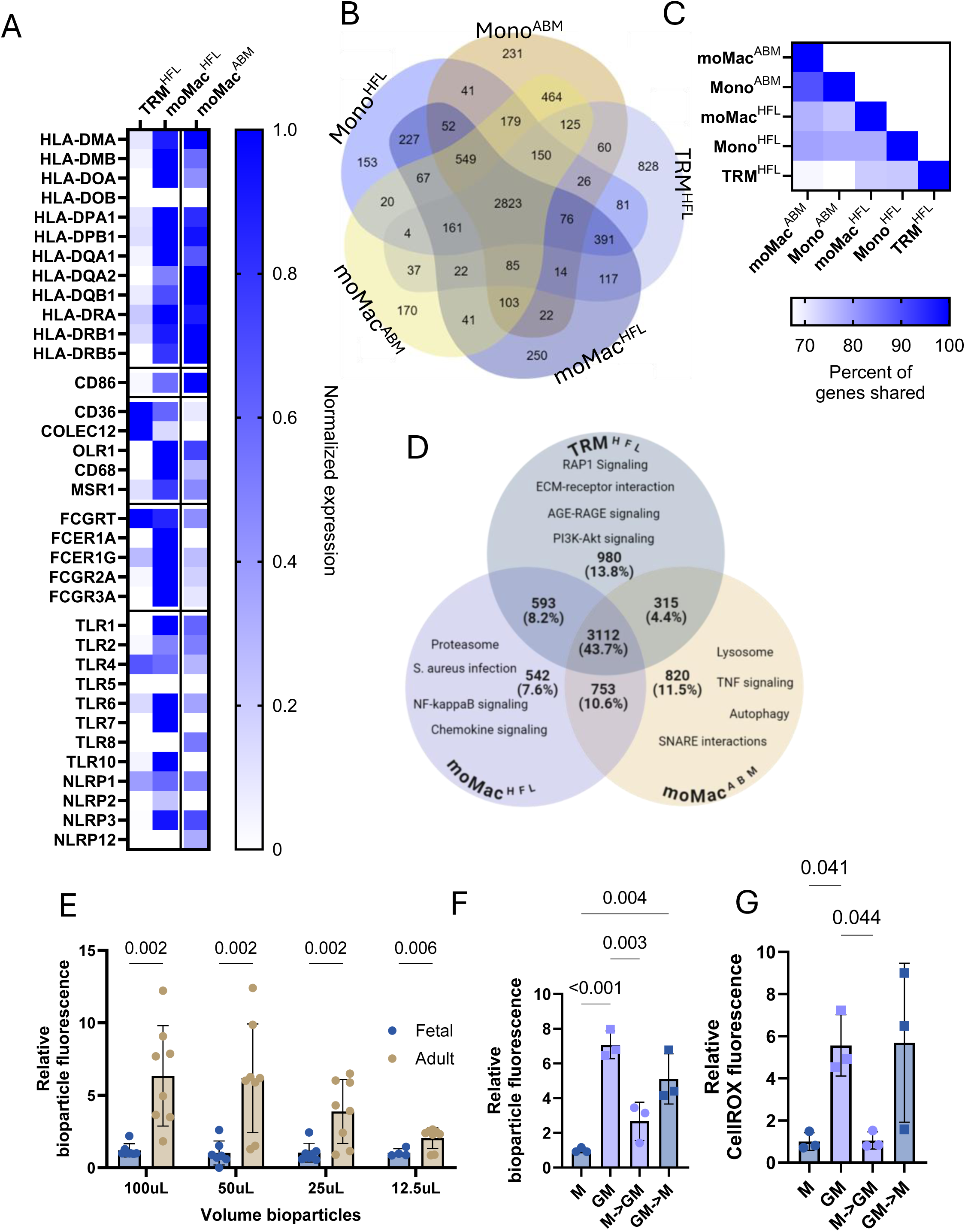
Expedited differentiation produces a macrophage with low inflammatory capacity. **A.** Heatmap of the indicated genes in TRM^HFL^, moMac^HFL^, and moMac^ABM^. Maximum expression for each gene is set to 1 and all others are normalized. **B.** Venn diagram of the 5000 highest expressed genes in Mono^HFL^, Mono^ABM^, TRM^HFL^, moMac^HFL^, and moMac^ABM^. **C.** Similarity index calculated from Figure 5B, where saturation indicated percent of genes shared between two populations. **D.** Venn diagram of the 5000 highest expressed genes amongst selected macrophages, annotated with the KEGG terms of highest significance for each individual macrophage. **E.** Phagocytic response of fetal and adult in vitro-differentiated macrophages ± SD. n = 8 for 25, 50, 100 µL doses; n = 4 for 12.5 µL dose. Multiple unpaired t-tests with two-stage step-up method. Data is presented as fold change from the average fetal response. Points represent biological replicates. **F.** Phagocytic response of fetal macrophages differentiated with either M-CSF or GM-CSF and then repolarized as indicated ± SD. n = 3; One-way ANOVA with Tukey’s multiple comparison’s test. **G.** ROS production post-phagocytosis measured by CellROX red fluorescence via flow cytometry. n = 3; One-way ANOVA with Tukey’s multiple comparison’s test.

### Retention of HSC-like self-renewal programs in TRMs

We hypothesized that the rapid change in cell state between CSF1R^+^ HFL HSCs and TRMs could provide a mechanism by which self-renewal potential is retained in TRMs, potentially regulated by *AHR*. Like the expression patterns in the scRNAseq, we observed lower overall expression of *AHR* in fMac^M-CSF^ compared to aMac^M-CSF^ **(Figure 6A)**. We hypothesized that this ligand-activated TF could be modulated as described in human HSPCs to also regulate macrophage self-renewal^71–73^. We therefore treated mature macrophages with StemRegenin 1 (SR1), an AHR antagonist, or FICZ, an AHR agonist^71^. We observed that at baseline, fMac^M-CSF^ showed prolonged proliferative potential compared to aMac^M-CSF^ in extended culture, consistent with baseline retained self-renewal **(Figure 6B)**. Upon SR1 treatment, fMac^M-CSF^ showed amplified expansion while adult and fetal moMacs did not show such a response **(Figure 6B-E)**. Agonism of AHR using FICZ appeared to have no effect in short term treatment but was a stressor in longer term treatment, and fMac^GM-CSF^ did not show a self-renewal response **(Figure 6C-E).**

**Figure 6:**
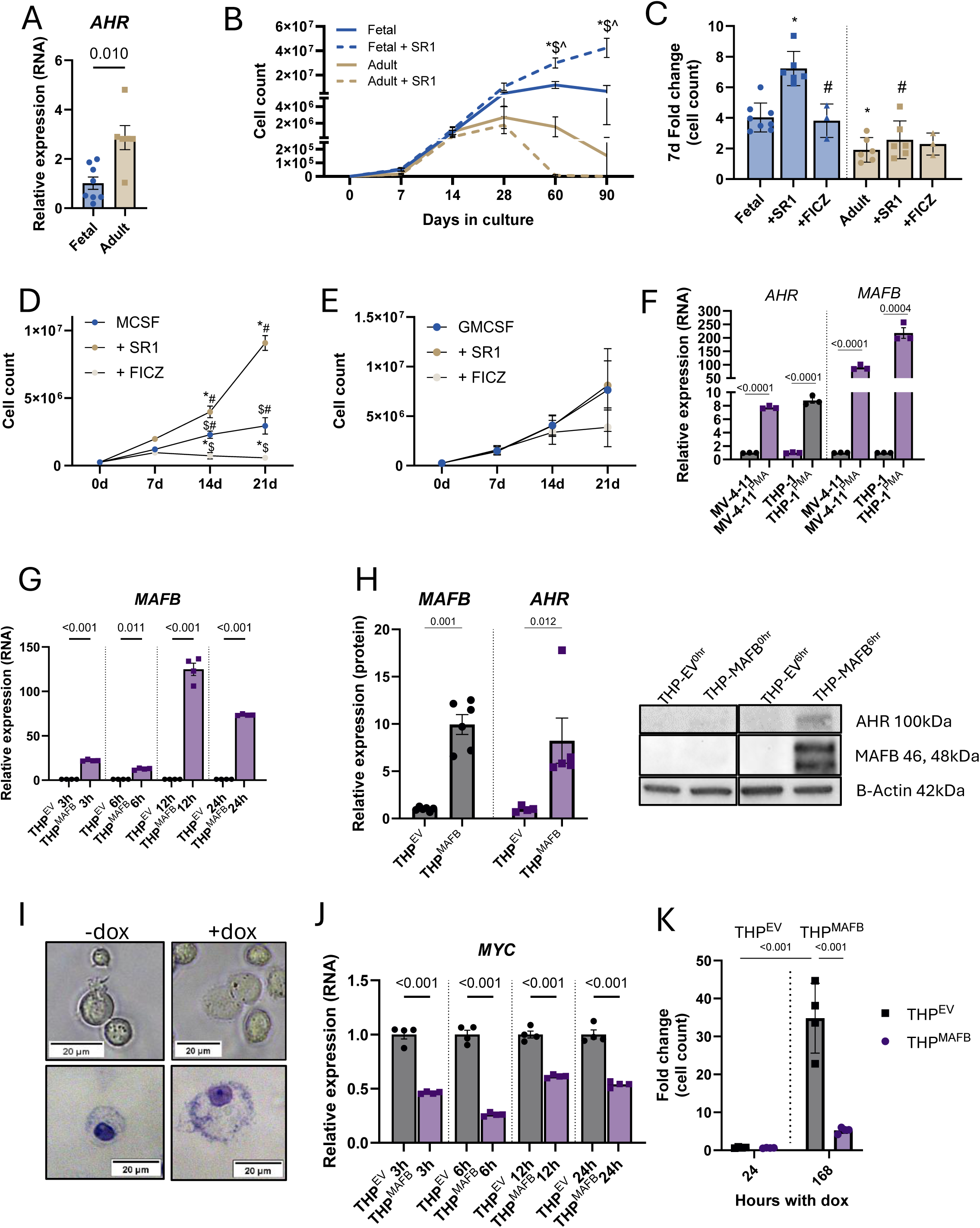
AHR modulation expands populations of directly differentiated macrophages. **A.** Expression of AHR in culture-differentiated fetal and adult macrophages, calculated as fold change from the average fetal expression. n = 6-8 biological replicates; each point is the average of at least 3 technical replicates. Unpaired two-tailed t-test. **B.** Number of macrophages in culture over time with and without treatment with 1 µM SR1, represented as cell count ± SEM. Treatment began after 14 days of M-CSF driven differentiation. 2-way ANOVA with Bonferroni’s correction; n = 4-5 biological donors per condition. * indicates p < 0.05 versus fetal, $ versus adult, and ^ versus adult + SR1. **C.** Effect of treatment with 1 µM SR1 or 1 µM FICZ on M-CSF-differentiated fetal and adult macrophages over 7 days. Cells were selected for CD14 prior to the start of the assay. n = 3-8 biological replicates per condition; one-way ANOVA with Tukey’s correction; * indicates p < 0.05 versus fetal, # versus fetal + SR1. **D.** Effect of treatment with 1 µM SR1 or 1 µM FICZ on fetal M-CSF- and **E.** GM-CSF-differentiated macrophages over 7, 14, and 21 days. Macrophages were selected for CD14 prior to the start of the assay. n = 3 biological replicates; two-way ANOVA with Sidak’s multiple comparisons test. **F.** Expression of AHR and MAFB in PMA-differentiated MV-4-11 and THP-1 cells measured by qPCR. Bars represent the fold change calculated from the average of the untreated controls ± SD. n = 3 individual experiments; two-way ANOVA with Bonferroni’s correction. **G.** Expression of MAFB in the doxycycline-inducible THP^MAFB^ cell line at the indicated time points following induction. Bars represent fold change from the average of the empty vector condition ± SD. n = 4 individual experiments; one-way ANOVA with Sidak’s multiple comparisons test. **H.** Protein expression of MAFB and AHR in THP^EV^ and THP^MAFB^ ± SD, expressed as fold change from THP^EV^, measured by western blot. n = 4 individual experiments. Mixed effects analysis with REML model and Sidak’s multiple comparisons test. Representative blots to the right. **I.** Images of uninduced THP^MAFB^ and induced THP^MAFB^ at 24 hours. Top: Phase contrast images of cells in culture. Bottom: Cytospins stained with Grunwald-Giemsa for visualization. Magnification 200x, scale bars represent 20 µm. **J.** Expression of MYC in the doxycycline-inducible THP^MAFB^ cell line at the indicated time points following induction, measured by qPCR. Bars represent fold change from the average of the empty vector (EV) condition ± SD. n = 4 individual experiments; one-way ANOVA with Sidak’s multiple comparisons test. **K.** Proliferation of THP^EV^ or THP^MAFB^ 24 hours and 168 hours treatment with doxycycline. Data is presented as fold change from a starting cell count of 10,000. n = 4 individual experiments; two-way ANOVA with Sidak’s multiple comparisons test.

MAFB has a known role in restricting myeloid and macrophage proliferation as well as directing monocyte to macrophage differentiation^32,64,74^. We observed high expression of both *MAFB* and *AHR* in aMac^M-CSF^ and speculated that MAFB could regulate the expression of *AHR* to enact its effects on proliferation, based on a predicted binding site of MAFB in the promoter region of AHR (human hg18 chr7:17,304,491-17,304,496) **(Supplemental Figure 6A)**^75,76^. This interaction could provide a potential mechanism for self-renewal pathways to integrate with core macrophage TF programs. We first found that inducing differentiation of THP-1 and MV-4-11 cells with phorbol 12-myristate 13-acetate (PMA) resulted in increased expression levels of both *MAFB* and *AHR* in the resulting macrophages **(Figure 6F)**. We then engineered a doxycycline-inducible *MAFB* THP-1 line that could trigger macrophage-like differentiation in THP-1 cells as early as 24 hours post-induction **(Figure 6G-I, Supplemental Figure 6B)**. Induction of *MAFB* led to increased AHR protein with concomitant downregulation of *MYC* **(Figure 6H-J)**. *MAFB* induction also greatly reduced THP-1 proliferation and produced a macrophage surface profile **(Figure 6K, Supplemental Figure 6C),** altogether suggesting that MAFB directs the expression of AHR, excluding terminal moMacs from the cell cycle.

### Selective TRM expansion mitigates inflammation in AD

Because we observed diminished inflammatory activity in TRMs and that TRMs – but not moMacs - could be expanded by AHR inhibition, we sought to leverage this effect therapeutically in a model of inflammation. In many inflammatory diseases, self-renewing TRMs die after initiation of inflammation and repopulate upon resolution^15^. We hypothesized that selective pharmacologic expansion of TRMs during acute inflammation could promote resolution. To verify the applicability of our main findings in mice, we first differentiated murine multipotent HSPCs using M-CSF, as with human HSPCs. We observed that murine fetal M-CSF-differentiated macrophages showed lower phagocytic capacity and ROS generation than adult macrophages, recapitulating observations in human cells **(Figure 7A-B)**. We next conducted in vitro testing with AHR antagonists SR1 and CH223191 on terminal mouse macrophages to confirm that the effects of AHR modulation were conserved between species. We observed that murine M-CSF-differentiated macrophages responded to AHR antagonism analogously to human cells **(Figure 7C)**. Next, to determine the therapeutic value of expanding Langerhans cells (LCs), the self-renewing TRMs of the epidermis, in a chronic inflammatory setting we employed two models of AD (**Supplemental Figure 7A)**. In the first model, after sensitization with dinitrochlorobenzene (DNCB), the mouse is treated on both ears with DNCB to induce AD. Each mouse was additionally treated with SR1 versus vehicle on contralateral ears **(Figure 7D)**. Skin thickening is a measure of severity in AD, corresponding to the influx of inflammatory immune cells, including moMacs, into the lesion. We observed SR1 treatment markedly reduced the gross severity of the disease as well as the thickness of the ear **(Figure 7E-G)**. Local lymph node mass was significantly reduced on the side of SR1 treatment **(Figure 7H).** We then assessed the macrophage content of the ears and lymph nodes and observed a reduced ratio of inflammatory Ly6c^high^ moMacs to F4/80^+^Ly6c^low/neg^ TRMs in SR1 versus vehicle-treated ears, which negatively correlated with ear thickness (**Figure 7I-J**). We repeated this experiment using the vitamin D3 model of AD in nude mice to isolate effects on the myeloid system with similar effects observed **(Supplemental Figure 7B-D).**

**Figure 7:**
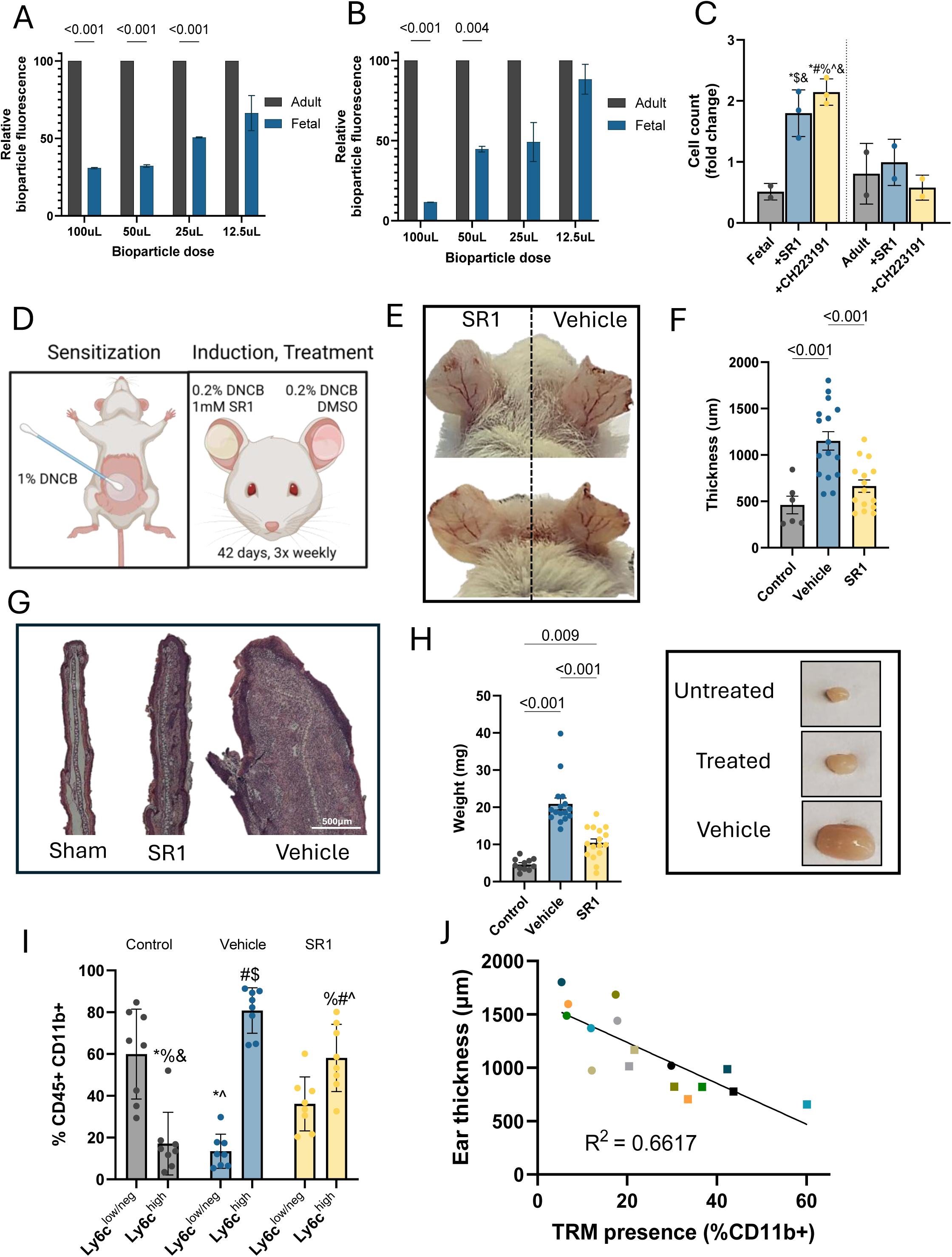
AHR modulation dampens inflammation and expands TRMs in AD models. **A.** Phagocytic response and **B.** ROS generated by murine M-CSF-differentiated fetal liver and BM macrophages. n=2-3 biological replicates. Multiple unpaired t-tests. **C.** Murine macrophage response to AHR modulation. Bars represent fold change in cell count after 7 days of treatment ± SD; n=2 biological replicates; one-way ANOVA with Tukey’s multiple comparisons test.; * indicates p < 0.05 vs fetal, # vs fetal + SR1, $ vs fetal + CH223191, % vs adult, ^ vs adult + SR1, and & vs adult + CH223191. **D**. Diagram depicting induction of AD with DNCB and treatment strategy. **E.** Image of mouse ears following 42 days of 3x weekly treatment with SR1 or vehicle. **F.** Ear thickness as a readout of AD severity. Mouse ear thickness was measured histologically and is reported in µm ± SD. n=6-16 biological replicates. Each point represents the average of at least 3 measurements. One-way ANOVA with Tukey’s multiple comparisons test. **G.** Images from histological sections of the outer edge of the ear. Scale bar represents 300µm; magnification 200x. **H.** Image and mass quantification ± SD of cervical lymph nodes from AD mice. Mice treated with SR1 had significantly lower lymph node mass, indicating lower inflammation. n=6-16 biological replicates. One-way ANOVA with Tukey’s multiple comparisons test. **I.** Quantification of Ly6c^++^ inflammatory moMacs and F4/80+ Ly6c−/low TRM in the ears of AD mice. Data is expressed as a percent of CD45^+^CD11b^+^ ± SD to exclude other myeloid populations. n=8-16 biological replicates; two-way ANOVA with Tukey’s multiple comparison’s test. **J.** Linear regression of paired ear thickness and TRM presence in the ears of AD mice. Colors correspond to individual mice; circles represent SR-1 treated ears; squares represent vehicle-treated ears.

## DISCUSSION

Over the past two decades, study of mononuclear phagocytes has elucidated macrophage heterogeneity, but the provenance of various macrophage subsets – particularly in the human - remains controversial. Though much is conserved between murine and human hematopoiesis, many of the paradigms of macrophage behavior established in mouse do not overlay directly onto human, making study of this compartment in human an unmet need due to the plethora of macrophage-driven diseases^77–79^. Studies in mice have compellingly demonstrated that some macrophages arise from primitive progenitors in utero^2,5,6,55,80–82^. However, the question of macrophage origin and specification in human largely relies on the assumption that human TRM development directly parallels that of the mouse. Given the differences in developmental pace, it is plausible that there are divergences in early macrophage generation. In this study, we focus on early human gestational ages (10-15 PCW) and demonstrate that a subset of definitive HSCs is specifically primed for macrophage differentiation, which occurs through an alternative, expedited pathway, resulting in a self-renewing macrophage with limited inflammatory capacity possessing hallmarks of a TRM state. This observation supports recent work demonstrating that HSCs are heterogeneous and that some HSC subsets are developmentally transient^31,83^.

### Ontogeny of TRMs

The precise identity of the cell giving rise to TRMs has been contested and use of varied lineage tracing systems has resulted in conflicting conclusions^84^. Reports of TRMs arising from pre-definitive erythro-myeloid progenitors (EMPs) from the yolk sac or fetal liver contrast other reports that TRMs arise from definitive hematopoietic cells^55,81,85^. EMPs are not well defined in humans and so conservation of these models remains unknown. It has also been reported in mice that a “pre-macrophage progenitor” arises in the fetal liver and seeds developing tissues simultaneously, where this cell differentiates and assumes a tissue-specific TRM identity^86^. We observe that a definitive macrophage-biased HSC expressing *CSF1R* exists in the HFL, differentiates rapidly in response to M-CSF, and seemingly bypasses a monocyte intermediate to generate TRMs. CSF1R is reportedly present on murine EMPs, while CD90 is present on definitive human fetal and adult HSCs, complicating identification of the equivalent murine cell^6,87^. In human, a “yolk sac-derived myeloid-biased progenitor” (YSMP), a possible equivalent of murine EMP, has been reported at Carnegie stage (CS) 11 (roughly 4 weeks PCW) but appears to be either absent or minimally present in both the yolk sac and HFL by CS17 (5-6 weeks PCW)^46^. YSMPs do not possess lymphoid potential akin to mouse EMPs, and also do not transcriptionally express CD90^46^. We employed the most robust selection possible, given the current body of knowledge of human pre-definitive progenitors: we restricted fetal donors to 10-15 PCW, a timepoint past reports of YSMP existence in the liver^46^. We sorted immunophenotypically definitive HSCs and observe that CSF1R-expressing HSCs show lymphoid potential and expression of genes marking definitive hematopoiesis^59,60^. We thereby conclude that TRM-biased HSCs are definitive in nature. This finding aligns with an independent report of developmentally-restricted expression of *CSF1R* in human fetal HSCs^60^. Further, YSMPs lack erythroid potential, which suggests interspecies divergence and highlights the need for detailed understanding of human developmental hematopoiesis^46^.

### Two available modes of differentiation utilizing distinct TFs

We observe two modes of macrophage differentiation available to HFL HSCs, which to our knowledge has not yet been reported. We capture the conventional model of cascading *SPI1*, *IRF8*, and *KLF4* expression with participation of other TFs to coordinate both fetal and adult monocyte fate^88–92^. However, expedited differentiation into self-renewing TRMs does not recruit the same TF repertoire. First, we observe two temporally distinct expression patterns for *SPI1* - it is moderately expressed in fetal and adult HSCs and MPPs, as would be expected due to its key role in maturation of the common myeloid progenitor as well as regulation of *CSF1R*, *CSF2R,* and *CSF3R*, encoding receptors for growth factors required for myeloid and monocytic maturation and survival^93–96^. Expression of *SPI1* diverges in more committed progenitors, increasing in expression in moMac^HFL^ and moMac^ABM^ differentiation, but decreasing in TRM^HFL^ differentiation. This is likely an important factor in commitment and maturation because of SPI1’s interactions with other important TFs for myeloid and monocytic differentiation, including CEBPA, CEBPB, IRF4 and IRF8^97–100^. IRF8 is indispensable for determination of granulocyte versus monocyte fate and coordinates with SPI1 to repress CEBP binding domains^101^. Level of IRF8 differentially affects myeloid lineage choice between granulocyte, monocyte, and dendritic cells in mice by shaping enhancer landscapes in early myeloid progenitors and promoting the expression of *KLF4*, which controls the transcription of terminal monocyte genes^88–92,101–103^. IRF8 is notably high in HSC^ABM^, but we do not observe this canonical pattern of expression of *IRF8* in TRM^HFL^ differentiation. We also observe attenuated expression of KLF4 in TRM^HFL^, suggesting that other TFs coordinate TRM specification.

We also observe a subset of HSCs expressing CSF1R, which allows them to directly respond to M-CSF signaling. In lieu of canonical differentiation via monocytes, these HSCs differentiate rapidly into macrophages in culture, apparently allowing these phagocytes to retain access to HSC-like programs governing cell cycle entry. *MAFB* overexpression in THP-1 cells was sufficient to induce macrophage differentiation, halt proliferation, and downregulate *MYC*, which mirrors the proliferative state after *MAF* and *MAFB* knockout in terminal macrophages^40^. TRM^HFL^ and fMac^M-CSF^ accordingly had high expression of *MYC* and low expression of *MAFB*, which permits these macrophages to proliferate. Since *CSF1R* activation does not ordinarily occur until late in monocytic differentiation as in the adult expression pattern, the presence of this receptor on a subset of HSCs likely impacts their fate determination and coordinates the expedited transition along the TRM^HFL^ differentiation trajectory. The result appears to be a macrophage that bypasses acquisition of inflammatory identity conferred by transit through the monocyte state, as evidenced by low expression of *HIF1A* and *NF-*κ*B*-related genes. Our scRNAseq expression data is supported by our in vitro assays that demonstrate transit through a monocyte intermediate permanently alters the macrophage’s responses. We observe that despite identical stimuli driving differentiation, fetal and adult macrophages have divergent immunophenotypes and phagocytic capacity, indicating that the environment is not the sole factor of specification of human macrophages. Further, our observations posit an explanation for how several distinct macrophage populations can coexist in a single tissue for long periods of time^36^.

### AHR as a regulator of macrophage self-renewal

AHR is a ligand-activated TF that acts as a regulator of HSC quiescence; it is dynamically downregulated upon exposure to growth factors to allow entry into the cell cycle^104^. This property has been leveraged to expand HSCs in culture by antagonism with small molecules, most commonly SR1^71,72^. We observe specific in vitro expansion of fMac^M-CSF^ – but not aMac^M-CSF^ - in response to SR1, suggesting that TRM self-renewal functions similarly to that of HSCs. These observations are further supported by profiling of human TRMs that exhibit high expression of self-renewal TFs *MYC*, *FLI1*, *MECOM*, *ERG*, and *ID1* compared to other macrophage populations. We do not observe this expression profile in HFL or ABM moMacs. AHR is characteristically high in monocytes, which we and others have observed^29,61^. This expression pattern explains the monocyte’s typical quiescence, which is notable because both the immediate upstream progenitors (e.g. GMPs and monoblasts) fetal-derived macrophages readily proliferate. This bottleneck in proliferative ability correlates with expression of *AHR* that peaks at the monocyte stage during differentiation (in moMac^HFL^ and moMac^ABM^). *MAFB* exhibits a similar pattern of expression, while there is a clear negative correlation with *MYC*. In THP-1 cells, we observe that *MAFB* induction stimulates differentiation while dynamically upregulating *AHR* and downregulating *MYC*, which then halts proliferation. This mechanism partly explains the observations that *MAFB* expression is anti-proliferative in myeloid cells and directly connects a core macrophage TF with blockade of self-renewal. The relatively low genetic similarity between TRMs and moMacs suggests that distinct AHR programs are engaged in these cells, with TRMs more closely resembling HSCs via their retained expression of self-renewal TFs. It also appears that expression of *AHR* is required for moMac survival, and while short term modulation is tolerated, longer term treatment has a detrimental effect on viability. Conversely, TRM survival does not rely on *AHR* or *MAFB* expression and it is likely that these cells dynamically adjust expression of *AHR* to permit re-entry into the cell cycle. Low expression of these TFs appears to allow high expression of *MYC* in TRM^HFL^ and cultured TRMs, consistent with feedback between these TFs.

### AHR as a therapeutic target

We observed a diminished inflammatory potential in TRMs and hypothesized that expanding TRMs in the midst of inflammation could promote resolution^105–109^. To query this, we used two models of AD, a chronic inflammatory condition characterized by an influx of immune cells, including moMacs^110–112^. LCs, the fetal-derived TRM of the epidermis, play an important role in initiation of inflammation in AD by signaling to circulating monocytes to carry out the acute inflammatory response and by transiting through the dermis to the draining lymph nodes to instruct T-cells^113^. Accordingly, LCs are almost absent in lesions with an influx of highly inflammatory moMacs^114^. By inducing AD and treating with SR1, we were able to shift the ratio of TRMs:moMacs toward homeostasis and reduce AD severity.

AHR has been proposed as a therapeutic target in glioma, chronic myeloid leukemia, and inflammatory diseases, and an AHR agonist is currently marketed for treatment of plaque psoriasis and AD^73,115,116^. In AD, AHR agonism reduces Th17 signaling, which alleviates inflammation^117^. We observe that agonism of AHR is detrimental to cell survival and speculate that we observe improvement in AD through a different mechanism, where agonism may be dampening proliferation of infiltrating inflammatory cells and antagonism expands TRMs.

### Limitations of the study

Human EMPs cannot be directly studied due to their proposed transient nature in early gestation and their existence in human is extrapolated from mouse studies, which limits our ability to define their contribution to human TRM pools. However, we verified the definitive nature of the TRM-primed HSC subpopulation based on multipotency, definitive *HOX* gene expression, and surface profile. Because TRMs appear to differentiate independently of canonical intermediates, it is possible that a TRM progenitor with an undefined immunophenotype exists in HFL or is so short-lived that it could not readily be captured. However, based on the rapidity of differentiation observed in vitro, we are skeptical of the existence of several progenitor intermediates between HFL HSCs and TRMs. We also interpreted differentiation history of macrophages (whether it transited through a monocyte intermediate) expression of transcripts expressed and inflammatory attributes acquired at the monocyte stage. However, macrophage plasticity could confound these parameters. We further acknowledge that macrophage potential is a dynamically regulated process that provides fine-tuned responses to environmental cues. In this study, we do not capture the dynamics of diseased microenvironments or macrophage reprogramming and instead focus mainly on cell intrinsic programs as a basis for further study of human macrophage ontogeny.

## ACKNOWLEDGEMENTS

RGR is supported by the NIDDK (R01DK134515). KF is supported by NHLBI (5T32HL007574-43). ELR is supported by Fundação de Âmparo à Pesquisa e Inovação de Santa Catarina (FAPESC, FAB2019121000026), and the Serrapilheira Institute (R-2111-39726) (ELR) We would also like to thank Ronald Mathieu, Mahnaz Paktinat, Ranjan Massey, Betelhem Gemechu, and Esha Budhiraja of the BCH-HSCI flow cytometry core for their expert assistance and valuable input.

## AUTHOR CONTRIBUTIONS

Conceptualization; KF, RGR

Methodology; KF, RGR, ELR

Investigation; KF, BB, DW, SF, CW, HK, HL

Analysis; KF, BB, ELDR, OM, HL, PC

Data curation; KF, RGR

Visualization; KF, OM, ELR

Writing; KF, RGR

Supervision; RGR

Resources; RGR, VGS

Funding; RGR

## SUPPLEMENTAL INFORMATION

Document S1. Supplemental figures 1-7

Table S1. Gene expression from scRNAseq in pseudobulk format. Excel file too large to fit in a PDF.

Table S2: List of genes used for cluster identification. Excel file too large to fit in a PDF.

Table S3. GO analysis of upregulated genes along each differentiation pathway. Excel file too large to fit in a PDF.

*Figure S1: Fetal HSPCs differentiate rapidly and proliferate in response to M-CSF*.

**A.** Gating strategy for sorting Lineage^−^CD34^+^CD38^−^ HSC/MPP. Lineage (CD3, CD14, CD16, CD19, CD20, CD56)-positive and dead cells were excluded by Pacific blue signal. Orange box indicates sorted population. Cells were sorted into 1.5mL tubes containing macrophage differentiation media. **B.** Scheme depicting macrophage differentiations. Banked CD34+ cells were sorted according to the gating strategy in A, then seeded into wells in the presence of SCF, Flt3L, TPO, M-CSF, IL-3, and IL-6. After 14 days, cytokines were withdrawn and macrophages were maintained with only M-CSF. Assays were conducted after 14-28 days in culture. **C.** Phase contrast image of a proliferative macrophage cluster in a fetal differentiation. 100x; scale bar = 100 µm. **D.** Images of cytospins of fMac^MCSF^ cultures, sampled daily. Labels indicate the number of days in culture with M-CSF. Magnification 200x; scale bars indicate 20µm. **E.** Flow cytometric measurement of surface markers of fMac^M-CSF^ and fMac^GM->M^. n=3 matched donors. Multiple paired t-tests with the Holm-Sidak method.

*Figure S2: HFL HSPCs produce TRMs in NSG-Q mice*.

**A.** Gating strategy for analysis of FRβ+ macrophage in HFL and ABM. Mononuclear cells were isolated by density gradient, then macrophages and monocytes were defined as live, CD14+. **B.** Human chimerism in the bone marrow of NSG-Q mice, expressed as a percent of total CD45+ cells ± SD. n=5 biological replicates per age; 2way ANOVA with Fisher’s uncorrected LSD. **C.** Human HSPCs, defined as CD34^+^, expressed as a percent of human CD45+ cells in the bone marrow of NSG-Q mice. n = 3 donors per age, unpaired, two-tailed t-test. **D.** Gating strategy for assessing human chimerism in bone marrow and tissues of NSG-Q mice. Single, live cells were gated as previously shown in S2A, then by expression of human or mouse CD45. **E.** Gating strategy for assessing human lineage composition in the bone marrow of NSG-Q mice. Single, live cells were gated as shown in S2A, then human cells by expression of human CD45. hCD45+ cells were then gated on expression of CD56 (NK cells), CD19 (B lymphocytes), CD3 (T-cells), CD14 (monocytes/macrophages, CD16 (granulocytes). **F.** Gating strategy for identifying FRβ+ macrophages in the liver, spleen, heart, and peritoneum of NSG-Q mice. Single, live cells were gated on expression of human CD45, then human CSF1R/CD115, then CD11b and FRβ. **G.** Total number of FRβ+ macrophages per organ of transplanted NSG-Q mice. n=2-5 donors per age; one-way ANOVA with Sidak’s multiple comparisons test. **H.** Peroxidase activity of human macrophages isolated from the livers of NSG-Q mice, as measured by resorufin fluorescence. n = 2 donors per age; unpaired, two-tailed t-test.

*Figure S3: scRNAseq populations and clustering*.

**A.** Target populations for single cell RNA sequencing and immunophenotypes for FACS isolation (top). Representative plots of sorted populations backgated onto FRβ and CD14. **B.** Gating strategy for the isolation of monoblasts, monocytes, and macrophages. Single cells were gated to exclude dead, CD3+, CD19+, and CD56+ cells by Pacific Blue signal. CD64+ cells were selected and monoblasts were isolated based on expression of HLADR without CD14. Monocytes and macrophages were defined as CD14, with CD163 and FRβ defining macrophages. Boxes indicate sorted populations. **C.** scRNAseq clusters before and after integration with Harmony. Light blue and blue indicate cells from individual donors. Clusters are indicated by individual colors.

*Figure S4: scRNAseq cluster descriptions*.

**A-B.** Signatures from the integrated clusters in S3C. **C.** Gene profiles of three types of macrophages found in the bone marrow, used to identify the type of macrophages in our scRNAseq dataset.

*Figure S5: CSF1R is expressed on a subset of definitive HFL HSCs*.

**A.** Expression of growth factor receptors on HSCs, MPPs, GMPs, and TRM intermediates in the present macrophage scRNAseq dataset. **B.** CSF1R expression in HFL HSCs from 10-14 PCW compared to expression in ABM HSCs (17-53 years) from the dataset of Li et al^31^. 23.3% of the fetal HSCs expressed CSF1R vs 4.2% of ABM HSCs. **C.** CSF1R expression in the ABM UMAP. **D-E.** Expression of transcription factors expressed in definitive HSCs in fetal progenitors and TRMs (C) and the across the lifespan (D)^31,59,60^.

*Figure S6: MAFB induction in THP-1 cells produces a macrophage-like cell*.

**A.** Consensus sequence of MAFB from Motifmap (cite). **B.** Phase contrast image of THP^MAFB^ cells 3 days post-induction of *MAFB,* exhibiting macrophage morphology. 200x, scale bar = 50µm. **C.** Flow cytometric analysis of THP^MAFB^ surface markers, normalized to EV control. N=4 individual experiments, two-way ANOVA with Holm-Sidak’s multiple comparisons text.

*Figure S7: Induction of eczema and Nu/J response to SR1 treatment*.

**A.** Schema for treatment schedules of the DNCB model of eczema (left) and the Vitamin D model of eczema (right). **B.** Lymph node mass of nude mice after induction of eczema, with and without treatment with SR1. One-way ANOVA with Sidak’s multiple comparison’s test. n=3-7 mice per condition. **C.** Measurement of ear thickness post-vitamin D induction of eczema. One-way ANOVA with Sidak’s multiple comparison’s test. n=3-7 mice per condition.

## STAR METHODS

### Data availability

*The sequencing data are available in the Gene Expression Omnibus (GEO) database at NCBI, accession number GSE284442, and can be accessed publicly at* https://www.ncbi.nlm.nih.gov/geo/query/acc.cgi?acc=GSE284442.

### Quantification and statistical analysis

Statistical analysis was performed in Graphpad Prism version 10. Analysis of flow cytometry was performed with FlowJo v10.10. Analysis of immunohistochemistry and Western blots was performed with Fiji ImageJ v2.15.1^118^. Details of statistical tests, statistical dispersion, number of biological donors, animals, or cells are reported in the figure legends. Significance was defined as p<0.05.

### Cells

Human bone marrow aspirates were obtained from healthy donors on a research protocol at Boston Children’s Hospital with the IRB approval number P00002781. Human fetal liver specimens were obtained through Advanced Bioscience Resources. Both female and male donors were obtained. Bone marrow donors were between 20 and 35 years old; gestational ages of human fetal liver donors were 10-15 PCW.

### CD34+ Isolation

Liver tissue was mechanically dissociated and incubated with 1mg/mL Collagenase IV (Gibco) in IMDM at 37° for 20 minutes, then passed through a blunt end syringe and a 70µm filter. Following dissociation of liver, the process of selection is identical for liver and marrow. Cell suspensions were diluted 1:1 with PBS and layered atop a Ficoll Premium Paque density gradient (Cytiva) and centrifuged for 40 minutes at 400xg at room temperature with the brake off. The interphase mononuclear layer was isolated and washed with PBS. CD34+ stem and progenitor cells were positively selected using magnetic microbeads for CD34 (Miltenyi) and cryopreserved at −196°C.

### Flow cytometry and sorting

CD34^+^ cells were thawed and incubated overnight for recovery in X-VIVO™-15 Serum-free Hematopoietic Cell Medium (Lonza) with supplemented Penicillin-Streptomycin (VWR2, cat. 45000-652), 50 ng/mL recombinant human SCF, Flt3L, TPO, IL6 and 10 ng/mL IL3 (Stemcell Technologies). The following day, cells were washed and resuspended in flow cytometry buffer (PBS +2%FBS +penicillin/streptomycin) and blocked with Trustain FcX (Biolegend) and Monocyte Block (Biolegend) for 10 minutes at room temperature. Following blocking, cells were stained for Lineage (anti-CD3, CD14, CD16, CD19, CD20, CD56, Pacific blue, Biolegend), CD34 (PE-Cy7, BD), and CD38 (PE, BD) at a dilution of 1µL/10^6^ cells for 20 minutes at room temperature. Cells were then filtered, washed, centrifuged at 300xg for 5 minutes, and resuspended in flow buffer containing 0.1µM Sytox Blue viability stain (Thermo) for sorting. Sorting of CSF1R-expressing HSCs followed an identical procedure with the addition of antibodies for CD90 (APC-Cy7, Biolegend) and CSF1R (PE, Biolegend) and a substitute CD38 antibody (FITC, Biolegend). Macrophage analysis followed an identical procedure using the antibodies FRβ (PE, Biolegend), CD64 (PE-Cy7, Biolegend), HLA-DR (FITC or PE-Cy7, Biolegend), CD86 (PerCP, Biolegend), CCR2 (APC-Cy7, Biolegend), CX3CR1 (FITC, Biolegend) and analyzed on an LSRII flow cytometer equipped with 355nm, 405nm, 488nm, 561nm, and 640nm lasers or a Cytek Aurora with 64 filters. Cell sorting was conducted on a BD Aria with 3 lasers or a BD Symphony with five lasers.

### Colony forming assay

Two thousand lineage^−^CD34^+^CD38^−^ cells were mixed 1.5mL MethoCult™ H4434 Classic (StemCell Technologies) containing fetal bovine serum, bovine serum albumin, SCF, IL3, EPO, and GM-CSF without antibiotics. The suspension was plated in a 35mm dish (Corning) and cultured in a humidified chamber. After 14 days, the colonies were blindly scored under a microscope.

### In vitro differentiation of macrophages

Lineage^−^CD34^+^CD38^+^ cells were sorted for macrophage differentiation assays. A similar procedure was followed for isolation of Lineage^−^CD34^+^CD38^−^CD90^+^CD115^+/−^ cells, with the addition of antibodies against CD90 (APC-Cy7, Biolegend) and CD115 (AF647, Biolegend). These cells were cultured in X-VIVO 15 (Lonza) with supplemented 50ng/mL recombinant human M-CSF, SCF, Flt3L, TPO, IL6 and 10 ng/mL IL3 (StemCell Technologies). After 10 days SCF, Flt3L, and TPO were withdrawn from the culture and macrophages were maintained in the presence of only M-CSF or GM-CSF.

### Selection of macrophages and treatment with AHR modulators

Experiments with AHR antagonists and agonists began with macrophage differentiation assays as described. Following 14 days of culture, macrophages were selected for CD14 using a biotinylated antibody (Biolegend) and streptavidin-labeled microbeads (Miltenyi) or by Mojosort CD14 negative selection (Biolegend) to separate mature macrophages from remaining progenitors in culture. Both processes utilized LS columns (Miltenyi) and a QuadroMACS™ Separator (Miltenyi) and followed manufacturer recommendations for concentration of microbeads and antibodies. Macrophages were then split into multiple wells and cultured in X-VIVO™-15 supplemented with 50ng/mL M-CSF or GM-CSF and 1 µM of SR1 (Selleck), CH-223191 (Selleck), or FICZ (MedChemExpress).

### Phagocytosis and ROS measurements

In vitro-differentiated macrophages were selected by CD14 microbeads as previously described. To isolate human macrophages from the organs of NSG-Q mice, we first digested the tissues as described for human fetal tissue. We then used a density gradient to isolate the human and mouse mononuclear fractions, which was then washed and selected for human CD14+ cells. 50,000 macrophages were evenly seeded in a 96 well format in Dulbecco’s Modified Eagle Medium (DMEM, Gibco) with the desired amount of bioparticles (0, 12.5, 25, 50, 75, 100 µL, Vybrant Phagocytosis assay, Thermo) to a volume of 100 µL. Following 1 hour of incubation at 37°C, 100uL of DMEM with 10 µM CellROX Red (Thermo) was added to each well (minus control wells) for a final concentration of 5 µM. For mitochondrial readouts, Mitotracker DeepRed (Thermo) and verapamil (Sigma) were added after 1 hour of incubation with final concentrations of 10 nM and 50 µM, respectively. Following an additional hour of incubation at 37°C, the cells were washed and prepared for flow cytometry.

### Peroxidase activity assay

Isolated macrophages were plated at a density of 50,000 per well in a 96 well format in X-VIVO 15 containing 20 mM H_2_O_2_ and 2 mM Amplex Red (Thermo) and allowed to incubate for two hours at 37°C. Resorufin fluorescence was recorded via CLARIOstar® High performance multimode microplate reader (BMG Labtech) at ex/em 570/585 nm with a gain of 1000.

### Culture of cell lines

THP-1 and MV-4-11 cells were regularly cultured in RPMI (Gibco) with 10% fetal bovine serum and 55 µM β-mercaptoethanol (Sigma) in T25 and T75 sized flasks (Thermo). Half the media was exchanged 2-3 times per week.

### Animal Handling and Husbandry

#### Mice

All animals were utilized in accordance with protocols approved by Boston Children’s Hospital Institutional Animal Care and Use Committees. Four NSG-Q (NOD.Cg-Prkdc^scid^ Il2rg^tm1Wjl^ Tg(CMV-IL3,CSF2,KITLG)1Eav Tg(CSF1)3Sz/J) mice were purchased from Jackson Laboratories (Strain #028657) to establish a breeding colony. Females were purchased with the genotype: Homozygous Prkdc scid, Hemizygous Il2rgtm1Wjl, Homozygous Tg(CMV-IL3,CSF2,KITLG)1Eav, Hemizygous Tg(CSF1)3Sz, and the males with the genotype Homozygous Prkdc scid Hemizygous Il2rgtm1Wjl, Homozygous Tg(CMV-IL3,CSF2,KITLG)1Eav, Non-Carrier Tg(CSF1)3Sz because homozygosity of human CSF1R renders the mouse sterile. Pups were genotyped for the CSF1 gene using the Jax-provided sequences and touchdown PCR cycle parameters using Taq polymerase (Takara). CSF1 primers; F: 5’-AAGGTGGGACAGTGA AGGTG-3’, R: 5’-GTGTGACACGTTGCCAAATC-3’. Four more female mice were purchased for experiments directly. We used female mice positive for CSF1 in transplants; for a total of 12. All other mice used were purchased directly for experiments.16 BALB/c were purchased from Charles River (Strain BALB/cAnNCrl). These mice were used in the DNCB model of eczema and were not bred in house. 12 Nu/Nu mice were purchased from Charles River (Strain Crl:NU-Foxn1nu). These mice were used in the Vitamin D3 model of eczema and were not bred in house. 6 NSG mice were purchased from Charles River for transplants of CSF1R^+/−^ HSCs. All mice used in transplants were female and between 10 and 14 weeks old at sacrifice by CO_2_ asphyxiation.

### Tail vein and bone marrow transplantation

Mice were preconditioned with 975 rad prior to transplants and subsequently maintained with antibiotics in drinking water (0.5 mg/ml trimethoprim/sulfamethoxazole). For transplants in NSG-Q mice, 1×10^5^ were injected into the tail vein. Mice were sacrificed 4 weeks post-transplant for analysis. In transplants of lineage^−^CD34^+^CD38^−^CD90^+^CSF1R^+/−^, mice were anesthetized with ketamine and 2000 cells in 10 µL sterile PBS were directly injected into the marrow of the femur. The marrow was aspirated at 4 weeks post-transplant and mice were sacrificed 12 weeks post-transplant for analysis.

### DNCB and vitamin D3 models of atopic dermatitis

We used two models of atopic dermatitis: 1-chloro-2,4-dinitrobenzene (DNCB, Sigma) and Vitamin D3 (VitD3, Merck). The DNCB-induced AD model is a 41-day treatment protocol broken into sensitization and induction phases, done in BALB/c mice^119^. At day 1, mice were sensitized with a single topical application of 1% DNCB (in 100% ethanol) to the abdomen using a cotton-tipped applicator following electric shaving with gentle restraint. The induction phase started on day 8 by application of 0.3% DNCB to both ears with a cotton-tipped swab, which was repeated three times a week through day 41. During this phase, we applied either vehicle (100% ethanol) and 1 mM SR1 to opposite ears. On day 42, mice were euthanized by carbon dioxide asphyxiation. The VitD3 model of atopic dermatitis requires application of 10 µL of 50 µM VitD3 (in 100% ethanol) to the front and back of each ear for four days, followed by a 3 day break, and a repeated four-day treatment period^111^. SR1 was applied consistently over the 11-day period to one ear, with the contralateral ear receiving only VitD3 or ethanol in the rest period.

### Processing of mouse tissue

All mice were euthanized by carbon dioxide asphyxiation. Bone marrow was obtained by isolating the mouse femur and tibia, removing the skin, muscle, and epiphysis, and flushing the inner compartment with PBS. The red blood cells were then lysed with red blood cell lysis buffer (Sigma), washed, and prepared for use in assays. Digestion of organs followed a similar protocol to isolation of CD34^+^ cells from human fetal tissue. Liver and spleen were minced with scissors and digested with 1mg/mL Collagenase IV (Gibco) in IMDM at 37° for 20 minutes, then passed through a blunt end syringe and a 70 µm filter. Heart, ear, and lymph node were finely chopped and digested with 0.2 mg/mL Liberase DH (Roche) in IMDM at 37° for 0.5-2 hours. Digested homogenates were then passed through a blunt ended syringe and a 70 µM strainer, washed, and prepared for assays. For certain assays, human cells were selected using either human CD45-biotin antibody (Biolegend) or negative CD14 selection (Biolegend) with streptavidin microbeads (Miltenyi).

### Protein isolation and western blotting

Protein was isolated from all protein sources using RIPA buffer (Pierce) with added protease inhibitors (Thermo) and brief sonication. 20µg of protein was diluted in Laemmli buffer and run on a 4-20% gradient gel (BioRad) for 60 minutes at 100 volts. Gels were transferred overnight onto PVDF membranes, blocked for two hours with 5% milk in TBS-T, and incubated with 1:2500 MAFB (Cell Signaling) and 1:2000 AHR (Cell Signaling) in 5% milk in TBS-T for 2 hours at room temperature. The membranes were washed three times and incubated with secondary antibodies at 1:5000 concentration for 1 hour.

### Quantitative rtPCR

Total RNA was extracted using TRIzol (Thermo) complementary DNA was synthesized with SuperScript II Reverse Transcriptase (Thermo). Quantitative PCR using Power SYBR™ Green PCR Master Mix (Thermo) and the QuantStudio™ 7 Flex Real-Time PCR System (Applied Biosystems) used the following cycling parameters: 95° C for 10 minutes cycles of 95° C for 15 seconds, 60° C for 60 seconds. The results were analyzed using the ddCT method. Human primer sequences were as follows: AHR; 5’-GTCGTCTAAGGTGTCTGCTGGA-3’, 5’-CGCAAACAAAGCCAACTGAGGTG-3’, GAPDH; 5’–GTCTCCTCTGACTTCAACAGCG-3’, 5’-ACCACCCTGTTGCTGTAGCCAA-3’, HES1; 5’-GGAAATGACAGTGAAGCACCTCC-3’, 5’-GAAGCGGGTCACCTCGTTCATG-3’, LYZ; 5’-ACTACAATGCTGGAGACAGAAGC-3’, 5’-GCACAAGCTACAGCATCAGCGA-3’, MAF; 5’- AGAAGTTGGTGAGCAGCGGCTT-3’, 5’-CACTGATGGCTCCAACTTGCGA-3’, MAFB; 5’- AGACGCCTACAAGGTCAAGTGC-3’, 5’-CGACTCACAGAAAGAACTCGGG-3’, MYC; 5’- CCTGGTGCTCCATGAGGAGAC-3’, 5’-CAGACTCTGACCTTTTGCCAGG-3’, S100A8; 5’- ATGCCGTCTACAGGGATGACCT-3’, 5’- AGA ATGAGGAACTCCTGGAAGTTA-3’

#### Lentiviral production and transduction

Tet-O-FUW-MAFB-P2A-mCherry was a gift from Andrea Califano (Addgene plasmid #203866), FUW-M2rtTA was a gift from Rudolf Jaenisch (Addgene plasmid #20342), FUW was a gift from David Baltimore (Addgene plasmid #14882), TFORF2469 was a gift from Feng Zhang (Addgene plasmid #142096), TFORF1577 was a gift from Feng Zhang (Addgene plasmid #143133), TFORF2950 was a gift from Feng Zhang (Addgene plasmid #144426)^120–123^. Plasmids were packaged with VSV-G (Addgene #12259) and PAX2 (Addgene #12260) and transfected into 293T cells using polyethylamine (Polysciences). Virus was collected at 24 and 48 hours and concentrated with Lenti-X™ Concentrator (Takara) and titrated using Lenti-X™ GoStix™ Plus (Takara). To generate the THP^MAFB^ line, THP cells were co-spinfected with Tet-O-FUW-MAFB-P2A-mCherry and FUW-M2rtTA, each at an MOI of 10, for 2 hours in a 24 well format. The viral media was removed, and the cells were allowed to recover for 7 days. The cells were then pulsed with doxycycline and flow sorted for mCherry positivity. Positive cells were sorted in bulk and as clones into conditioned media and cultured as normal without doxycycline.

### 10X Genomics single cell RNA sequencing

#### Isolation of cell populations

Unfractionated human fetal liver and adult bone marrow mononuclear fractions (Lonza) were thawed and CD34+ cells were selected using microbeads as previously described. CD34^+^ cells were then stained for Lineage (Pacific Blue CD3, CD14, CD16, CD19, CD20, CD56, Biolegend #348805), CD34 (PECy™-7 BD #348791), CD38 (PE, BD# 555460) and resuspended in flow buffer containing 0.1µM Sytox Blue viability stain (Thermo). From this subset of cells, live lineage^−^CD34^+^CD38^−^ and lineage^−^CD34^+^CD38^+^ cells were sorted and included in downstream analyses. Cells were sorted on a BD Fusion Aria with 405nm, 488nm, 561nm, and 640nm lasers.

CD34^−^ cells were stained for human CD19, CD3, CD56 (all Pacific Blue™, Biolegend), CD64 (PECy™-7, Biolegend), CD14 (BV605, Biolegend), CD11b (AF700, Biolegend), CCR2 (APCCy™-7, Biolegend) HLA-DR (FITC, Biolegend), CD163 (PerCP, Biolegend) and FOLRβ (PE, Biolegend). Dead, CD19^+^, CD3^+^, and CD56^+^ cells were excluded by Pacific Blue positivity and the remaining cells were gated on CD64^+^. CD14^−^ HLADR^++^ (monoblasts), CD14^+^HLADR^+^FOLR2^−^ (monocytes) and CD14^+^HLADR^+^FOLR2^+^(macrophages) were sorted using a BD Aria with 355 nm, 405 nm, 488 nm, 561 nm, and 640 nm lasers.

### Preparation of libraries

After sorting, cells were counted, fixed, and permeabilized. Libraries were comprised of 12,000 cells; 2000 lineage^−^CD34^+^CD38^−^, 2000 lineage^−^CD34^+^CD38^+^, 2000 monoblasts, 2000 monocytes, and 4000 macrophages. Tagmentation, GEM generation, and barcoding were carried out according to manufacturer’s instructions using reagents included in the Single Cell Multiome ATAC + Gene Expression kit (10X Genomics). Pre-amplification was carried out using NEBNext® High Fidelity (New England Biolabs). The rest of the preparation was carried out according to 10X recommendations in User Guide CG000338 (10X Genomics).

### Sequencing

Libraries were pooled and sequenced in two lanes on a NovaSeq S4 at a depth of 20,000 reads per cell for the RNA library.

### Analysis of single-cell RNA-sequencing data

Samples were divided according to stage, adult and fetal; each sample group had 2 replicates. Count matrices of genes × cells from each sample were imported in the R v. 4.3.0 statistical environment and processed into individual Seurat objects (Seurat v.5.1, which was used as the primary analytical package). Data processing, integration, normalization and every other analysis were performed independently on each sample group, using the same parameters. Normalization to 10,000 transcripts per cell barcode was run on each sample individually (log-transformed (NormalizeData)), and variable features were identified using the vst method based on the top 2,000 features (FindVariableGenes).

Cell cycle correction was performed to preserve the differentiation signal. Each cell within the dataset was given continuous cell cycle scores and assigned a discrete cell cycle state (CellCycleScoring), followed by the regression of the difference between the S and G2M scores. Individual replicates were then integrated using Seurat’s IntegrateLayers procedure, using 2,000 anchors, 30 dimensions and HarmonyIntegration as the chosen method. After the integration of each replicate, UMAP, kNN and clustering methods were calculated on the integrated reduced dimensions. The top 30 PCs were used to build the kNN graph, considering the 20 nearest neighbors. The resulting graph was partitioned using the Louvain shared nearest neighbor (SNN) modularity optimization-based clustering algorithm at resolution 0.5 to identify clusters (FindClusters). The cluster-specific gene markers were identified from the integrated, normalized gene expression data using a logistic regression test, 0.25 for both logfc.threshold and min.pct parameters, and returning only positive markers (FindAllMarkers). The clusters were then manually examined and annotated based on hematopoietic lineage gene markers^124^.

Trajectory analysis was performed using CellRouter by defining initial and final cell states, as previously described^48,125^.

## KEY RESOURCES TABLE

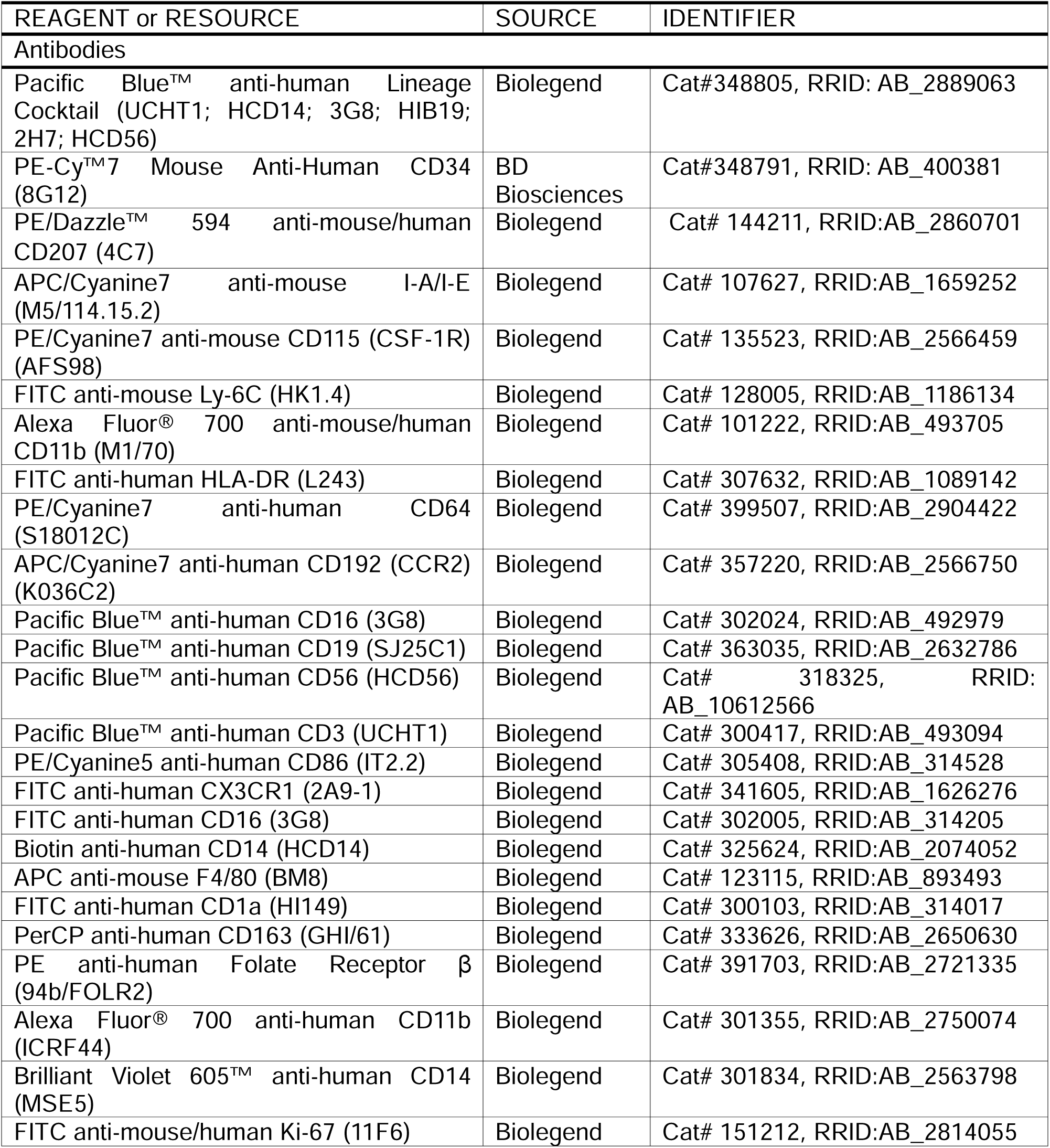
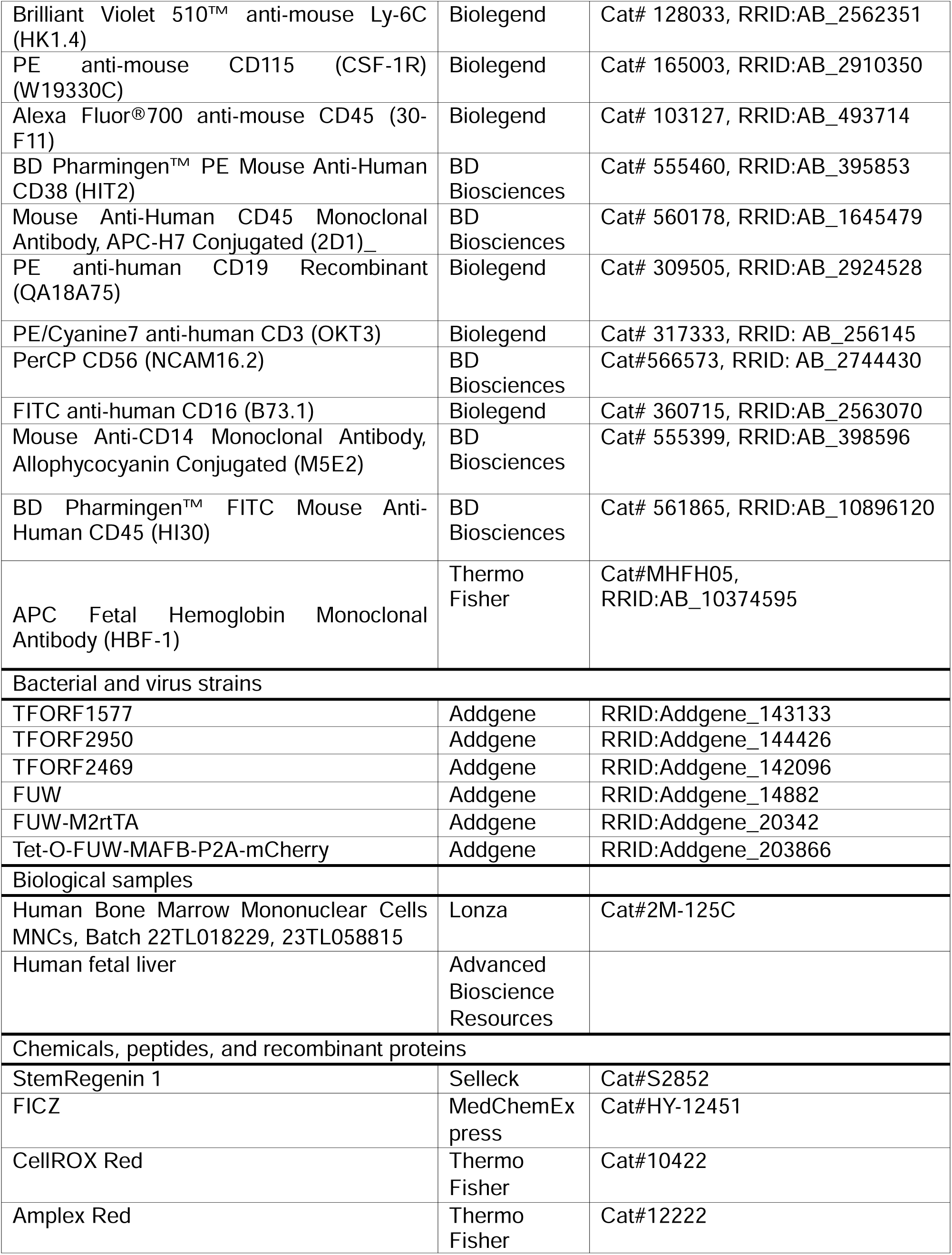
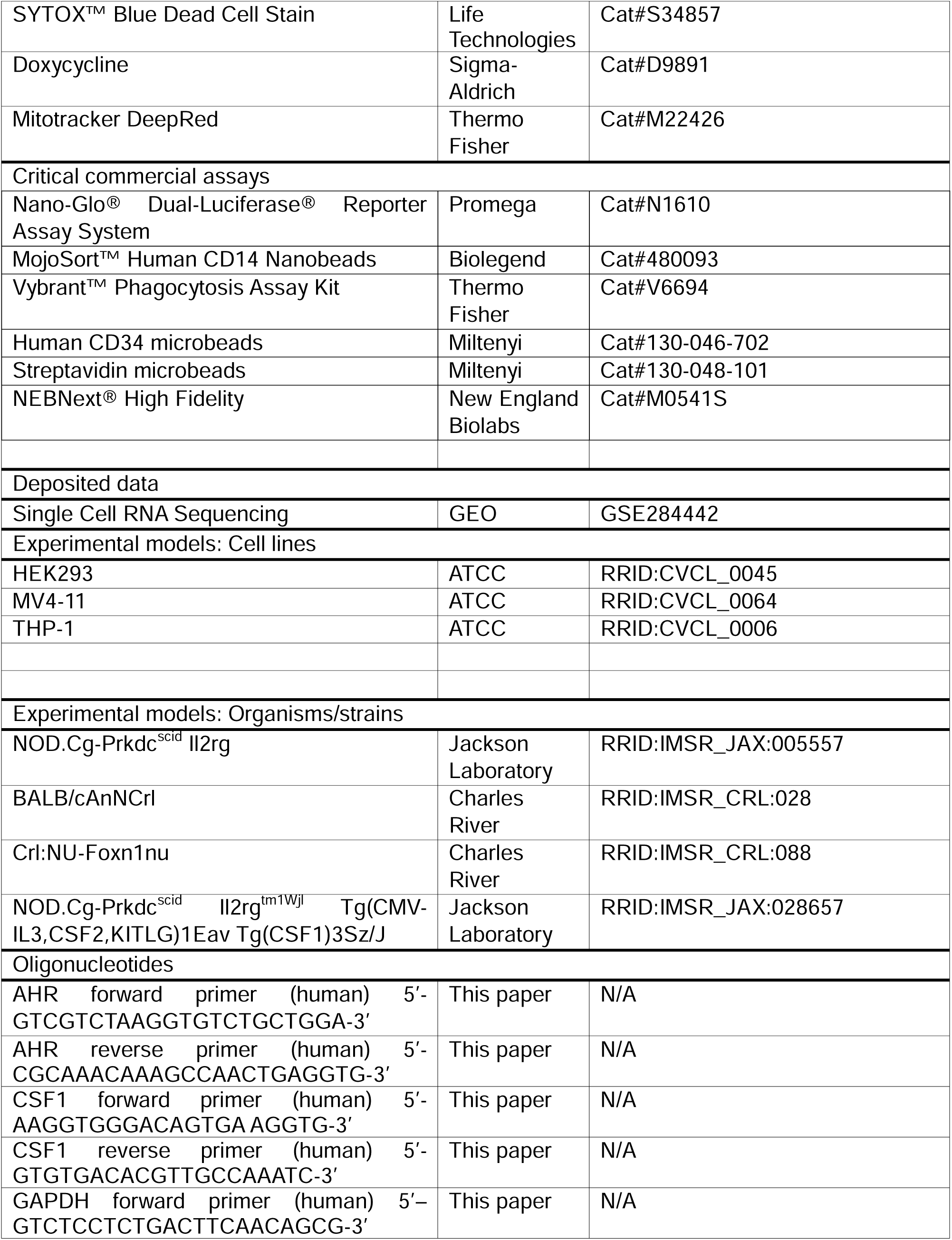
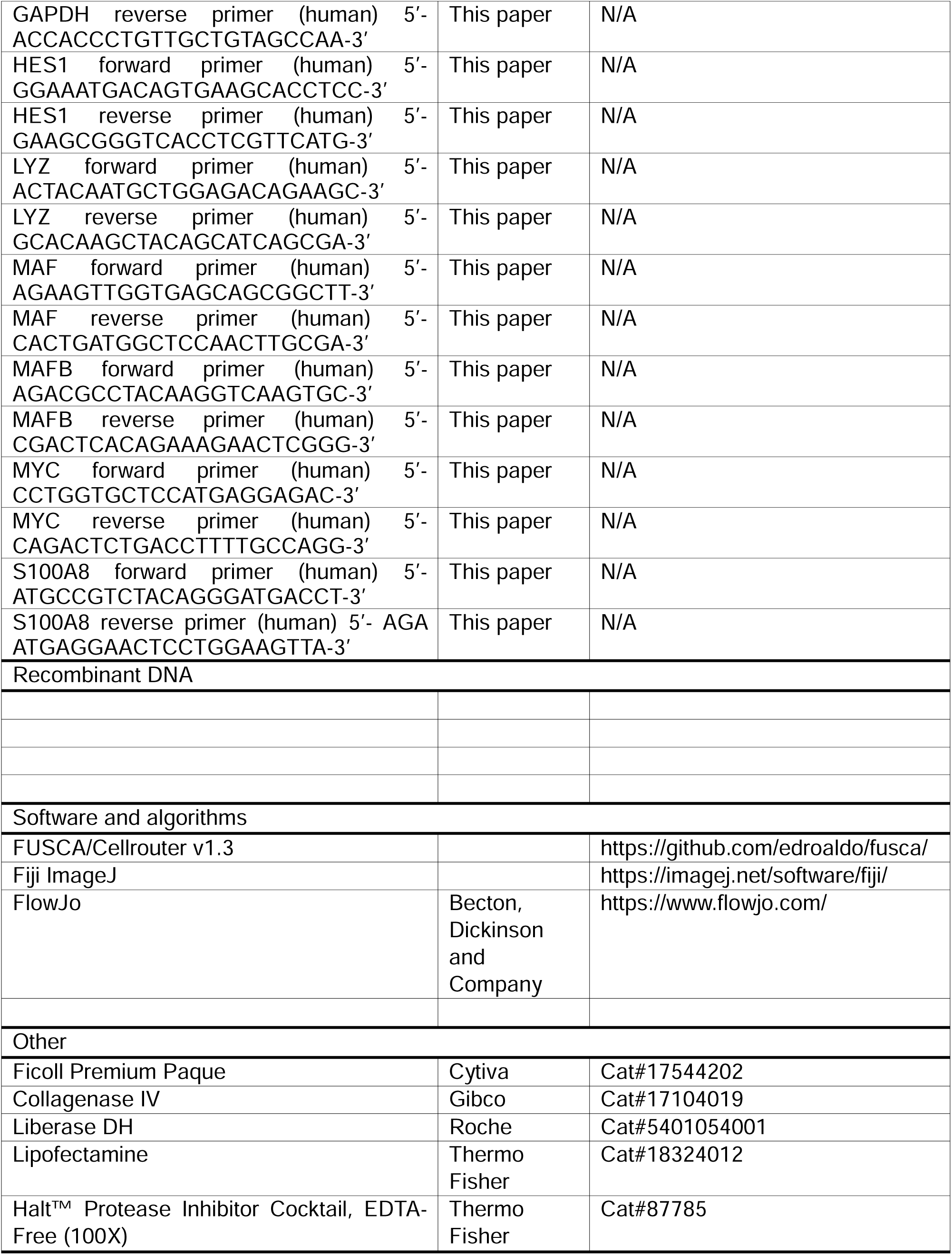

## LIST OF ABBREVIATIONS

ABM: Adult bone marrow
AD: Atopic dermatitis
AHR: Aryl hydrocarbon receptor
CFU: Colony forming assay
CS: Carnegie stage
EMP: Erythromyeloid progenitor
EV: Empty vector
FRβ: Folate receptor beta
GO: Gene ontology
GM-CSF: Granulocyte-macrophage colony stimulating factor
GMP: Granulocyte-monocyte progenitor
HFL: Human fetal liver
HSC: Hematopoietic stem cell
HSPC: Hematopoietic stem and progenitor cells
IL: Interleukin
LC: Langerhans cell
M-CSF: Macrophage colony stimulating factor
MFI: Mean fluorescent intensity
MPP: Multipotent progenitor
moMac: Monocyte-derived macrophage
PAMP: Pathogen-associated molecular pattern
PCW: Post-conception week
ROS: Reactive oxygen species
scRNAseq: single cell RNA sequencing
SR1: StemRegenin1
TF: Transcription factor
TRM: Tissue resident macrophage
YSMP: Yolk sac-derived myeloid-biased progenitor

