## Supplemental Figures for "Path of differentiation defines human macrophage identity"

### Figure S1

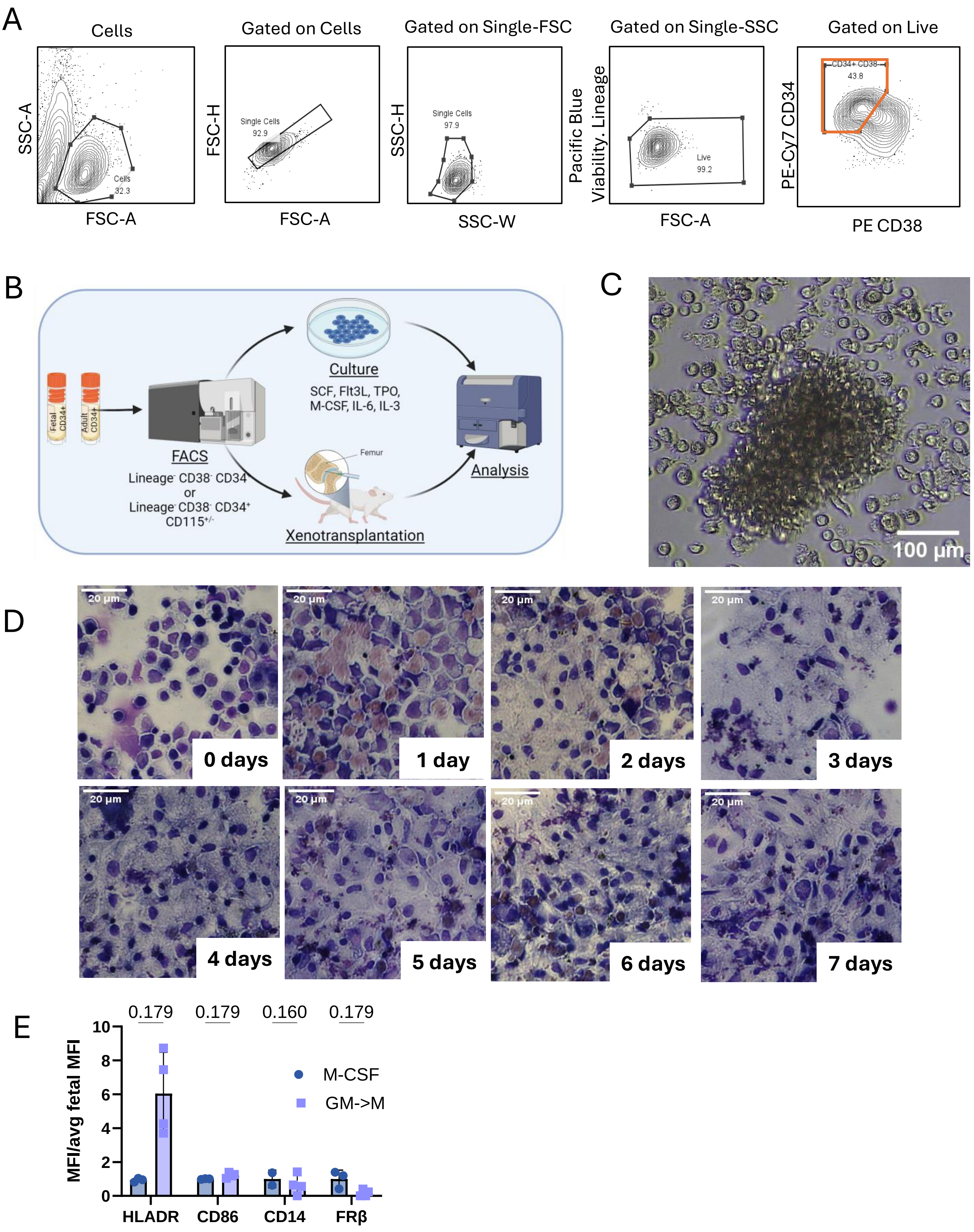
A
Cells
SSC-A
FSC-A
Cells
32.3
Gated on Cells
FSC-H
FSC-A
Single Cells
92.9
Gated on Single-FSC
SSC-H
SSC-W
Single Cells
97.9
Gated on Single-SSC
Pacific Blue
Viability. Lineage
FSC-A
Live
99.2
Gated on Live
PE-Cy7 CD34
PE CD38
CD34+ CD38-
43.8
B
Fetal CD34+
Adult CD34+
FACS
Lineage- CD38- CD34+
or
Lineage- CD38- CD34+ CD115+/-
Culture
SCF, Flt3L, TPO, M-CSF, IL-6, IL-3
Femur
Xenotransplantation
Analysis
C
100 µm
D
20 µm
0 days
1 day
2 days
3 days
4 days
5 days
6 days
7 days
E
MFI/avg fetal MFI
0.179
0.179
0.160
0.179
M-CSF
GM->M
HLADR
CD86
CD14
FRβ

### Figure S2

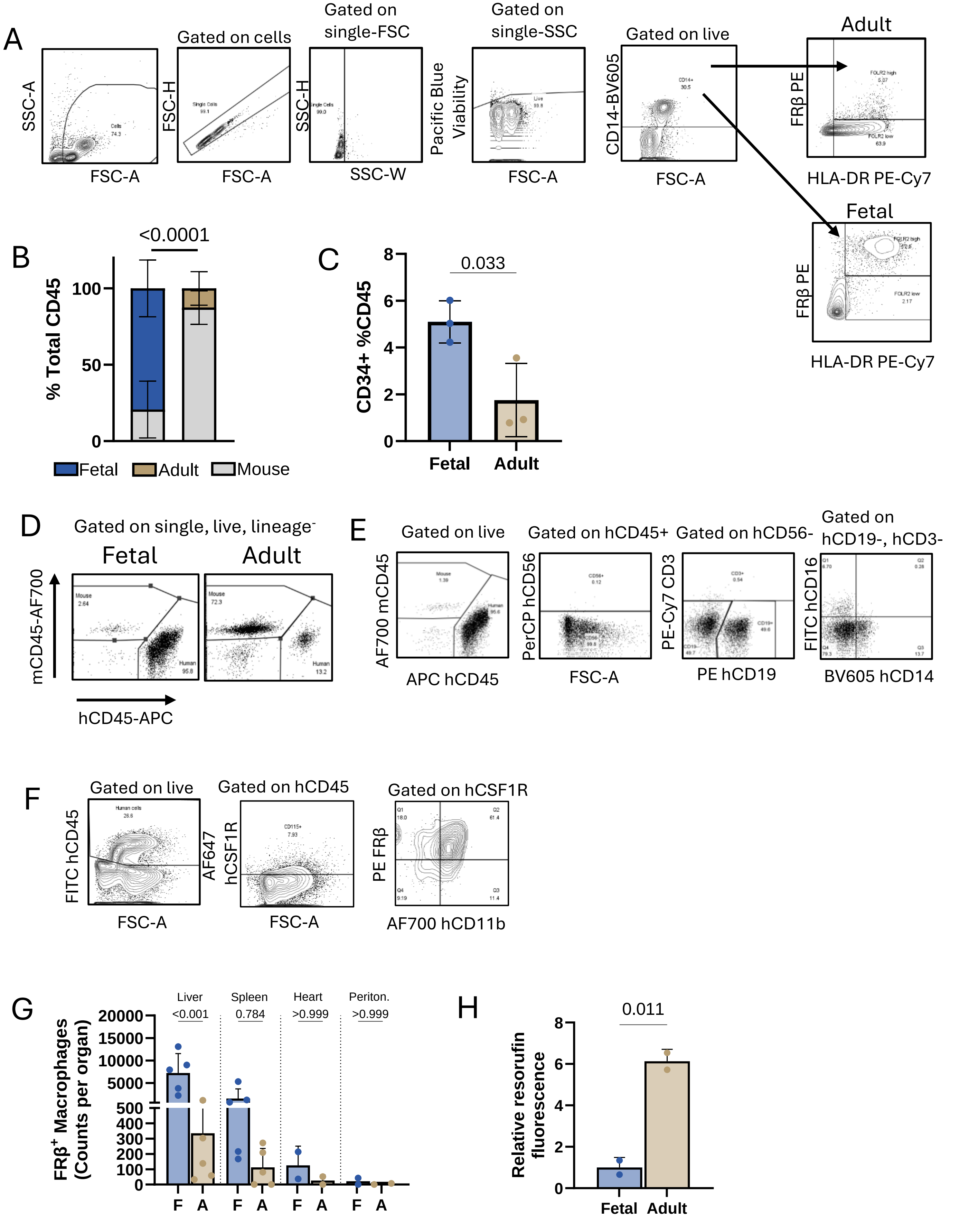
A
Gated on cells
Gated on single-FSC
Gated on single-SSC
Gated on live
Adult
Fetal
SSC-A
FSC-A
FSC-H
SSC-H
SSC-W
Pacific Blue Viability
CD14-BV605
FRβ PE
HLA-DR PE-Cy7
B
<0.0001
% Total CD45
Fetal
Adult
Mouse
C
0.033
CD34+ %CD45
Fetal
Adult
D
Gated on single, live, lineage-
Fetal
Adult
mCD45-AF700
hCD45-APC
E
Gated on live
Gated on hCD45+
Gated on hCD56-
Gated on hCD19-, hCD3-
AF700 mCD45
APC hCD45
PerCP hCD56
FSC-A
PE-Cy7 CD3
PE hCD19
FITC hCD16
BV605 hCD14
F
Gated on live
Gated on hCD45
Gated on hCSF1R
FITC hCD45
FSC-A
AF647 hCSF1R
PE FRβ
AF700 hCD11b
G
Liver
Spleen
Heart
Periton.
<0.001
0.784
>0.999
>0.999
FRβ+ Macrophages (Counts per organ)
F
A
H
0.011
Relative resorufin fluorescence
Fetal
Adult

### Figure S3

A

| Immunophenotype | Population |
|---|---|
| CD45+ Lin-CD34+CD38- | Multipotent HSPCs |
| CD45+ Lin-CD34+CD38+ | Progenitors |
| CD45+ CD34- CD19- CD64+ HLADR+ | Monoblasts |
| CD45+ CD64+ CD11b+ CD14+ CD163-/FOLR2- | Monocytes |
| CD45+ CD64+ CD11b+ CD14+ CD163+/FOLR2+ | Macrophages |

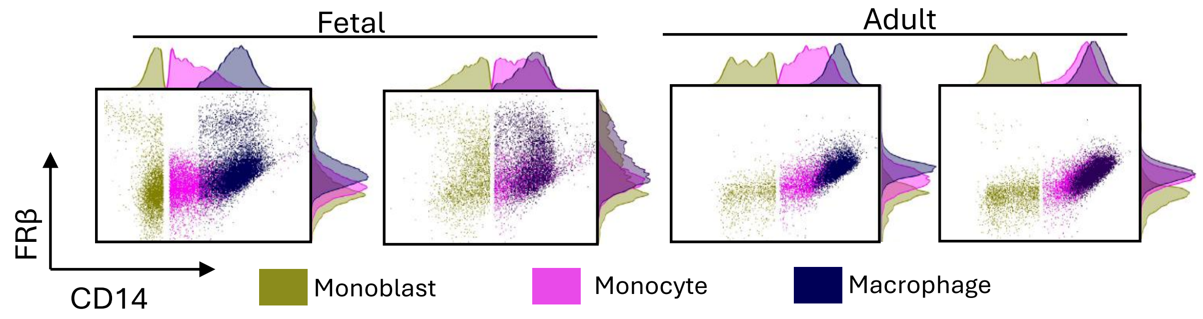


B

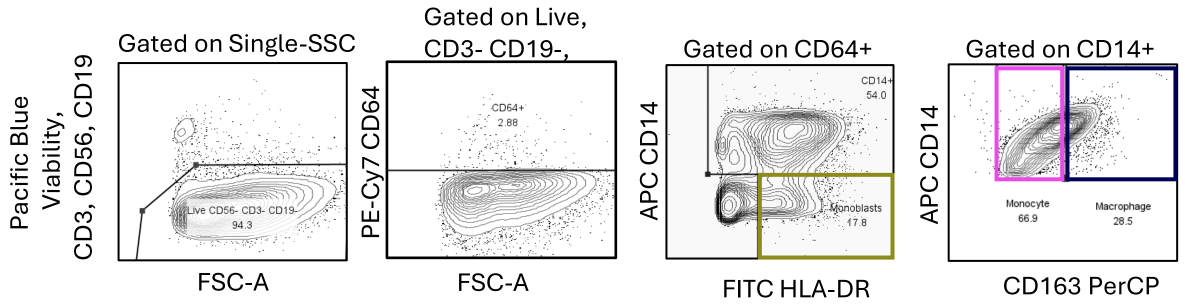


C

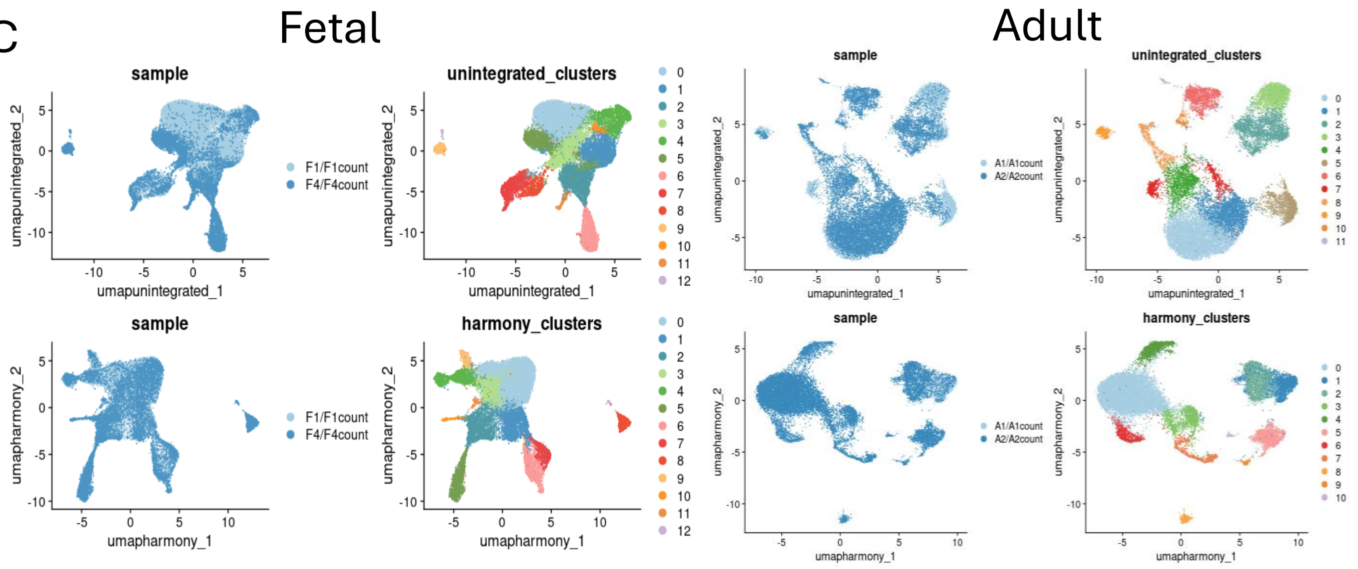

### Figure S4

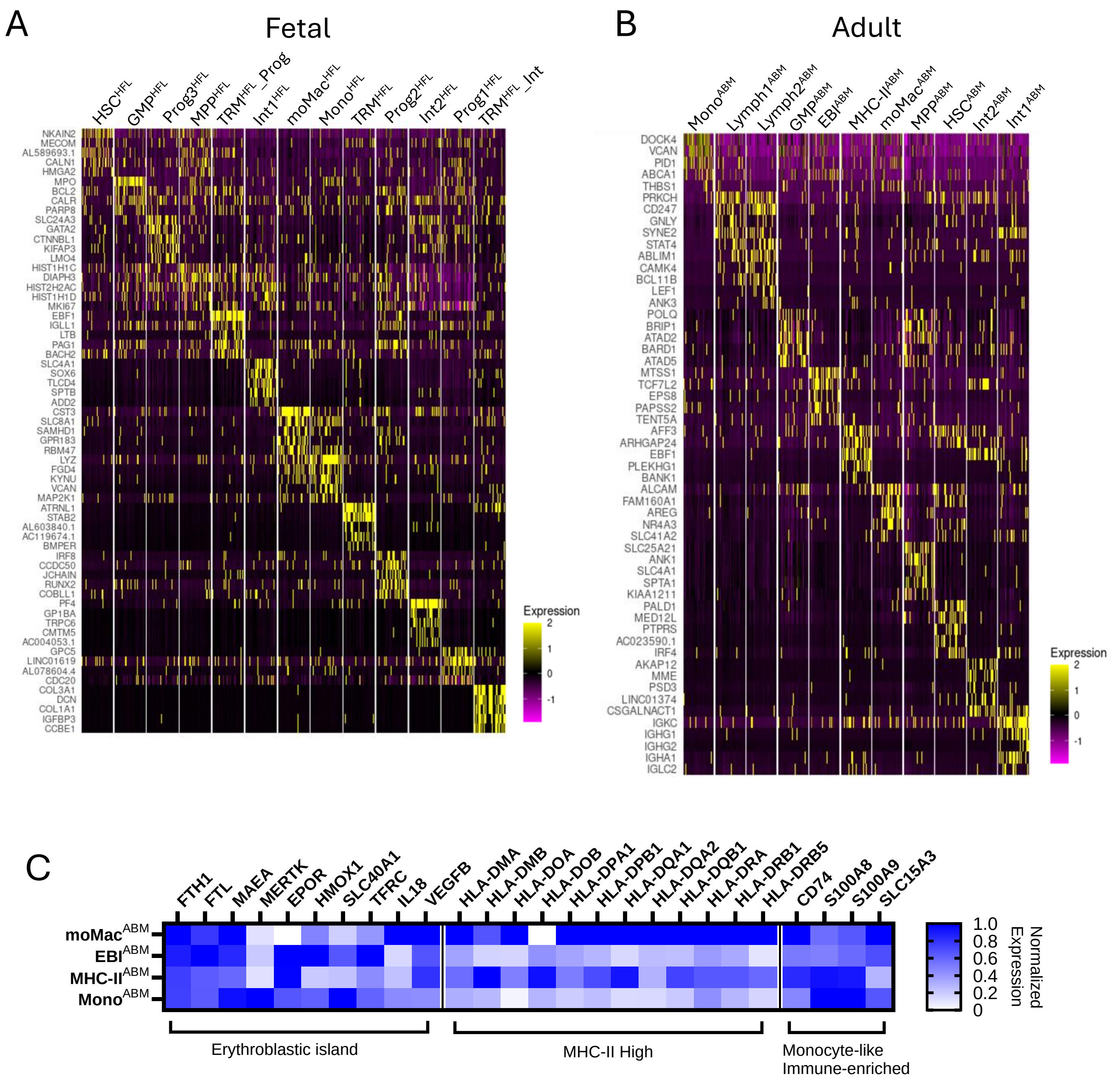

A
Fetal
HSC$^{HFL}$
GMP$^{HFL}$
Prog3$^{HFL}$
MPP$^{HFL}$
TRM$^{HFL}$_Prog
Int1$^{HFL}$
moMac$^{HFL}$
Mono$^{HFL}$
TRM$^{HFL}$
Prog2$^{HFL}$
Int2$^{HFL}$
Prog1$^{HFL}$
TRM$^{HFL}$_Int
NKAIN2
MECOM
AL589693.1
CALN1
HMGA2
MPO
BCL2
CALR
PARP8
SLC24A3
GATA2
CTNNBL1
KIFAP3
LMO4
HIST1H1C
DIAPH3
HIST2H2AC
HIST1H1D
MKI67
EBF1
IGLL1
LTB
PAG1
BACH2
SLC4A1
SOX6
TLCD4
SPTB
ADD2
CST3
SLC8A1
SAMHD1
GPR183
RBM47
LYZ
FGD4
KYNU
VCAN
MAP2K1
ATRNL1
STAB2
AL603840.1
AC119674.1
BMPER
IRF8
CCDC50
JCHAIN
RUNX2
COBLL1
PF4
GP1BA
TRPC6
CMTM5
AC004053.1
GPC5
LINC01619
AL078604.4
CDC20
COL3A1
DCN
COL1A1
IGFBP3
CCBE1
Expression
2
1
0
-1
B
Adult
Mono$^{ABM}$
Lymph1$^{ABM}$
Lymph2$^{ABM}$
GMP$^{ABM}$
EBI$^{ABM}$
MHC-II$^{ABM}$
moMac$^{ABM}$
MPP$^{ABM}$
HSC$^{ABM}$
Int2$^{ABM}$
Int1$^{ABM}$
DOCK4
VCAN
PID1
ABCA1
THBS1
PRKCH
CD247
GNLY
SYNE2
STAT4
ABLIM1
CAMK4
BCL11B
LEF1
ANK3
POLQ
BRIP1
ATAD2
BARD1
ATAD5
MTSS1
TCF7L2
EPS8
PAPSS2
TENT5A
AFF3
ARHGAP24
EBF1
PLEKHG1
BANK1
ALCAM
FAM160A1
AREG
NR4A3
SLC41A2
SLC25A21
ANK1
SLC4A1
SPTA1
KIAA1211
PALD1
MED12L
PTPRS
AC023590.1
IRF4
AKAP12
MME
PSD3
LINC01374
CSGALNACT1
IGKC
IGHG1
IGHG2
IGHA1
IGLC2
Expression
2
1
0
-1
C
FTH1
FTL
MAEA
MERTK
EPOR
HMOX1
SLC40A1
TFRC
IL18
VEGFB
HLA-DMA
HLA-DMB
HLA-DOA
HLA-DOB
HLA-DPA1
HLA-DPB1
HLA-DQA1
HLA-DQA2
HLA-DQB1
HLA-DRA
HLA-DRB1
HLA-DRB5
CD74
S100A8
S100A9
SLC15A3
moMac$^{ABM}$
EBI$^{ABM}$
MHC-II$^{ABM}$
Mono$^{ABM}$
Erythroblastic island
MHC-II High
Monocyte-like
Immune-enriched
1.0
0.8
0.6
0.4
0.2
0
Normalized
Expression

Supplemental Figure S5

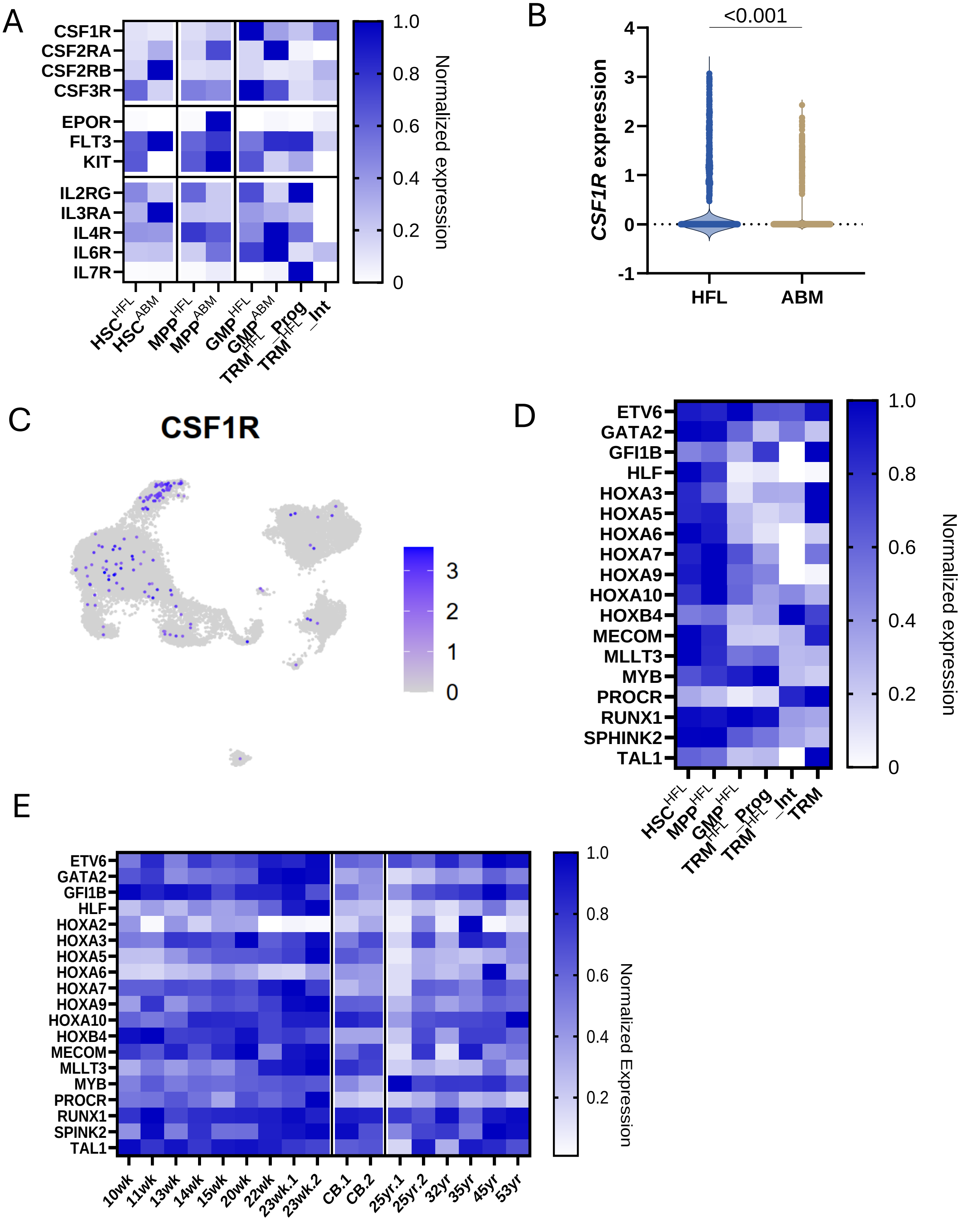

### Figure S6

A

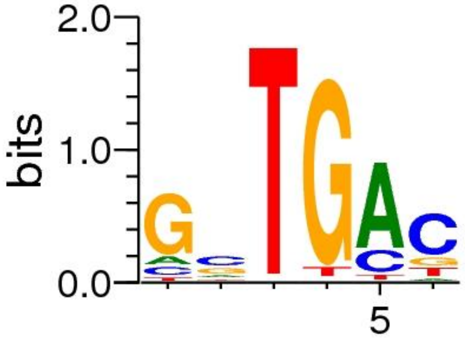


B

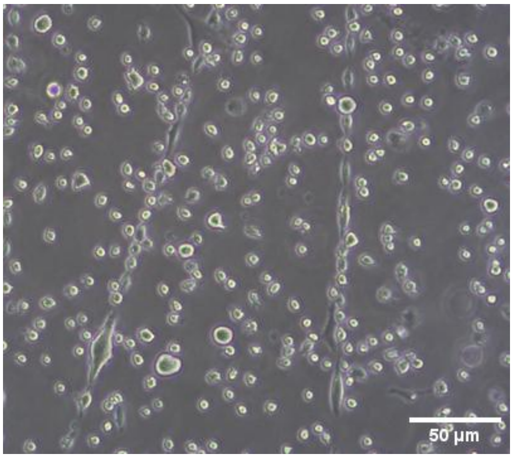


C

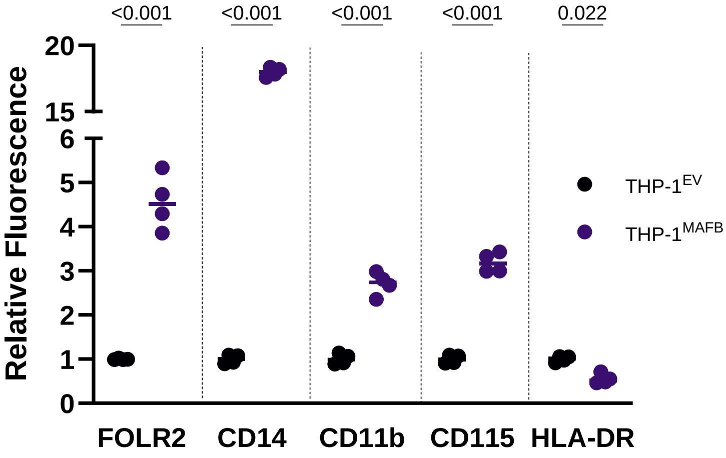

### Figure S7

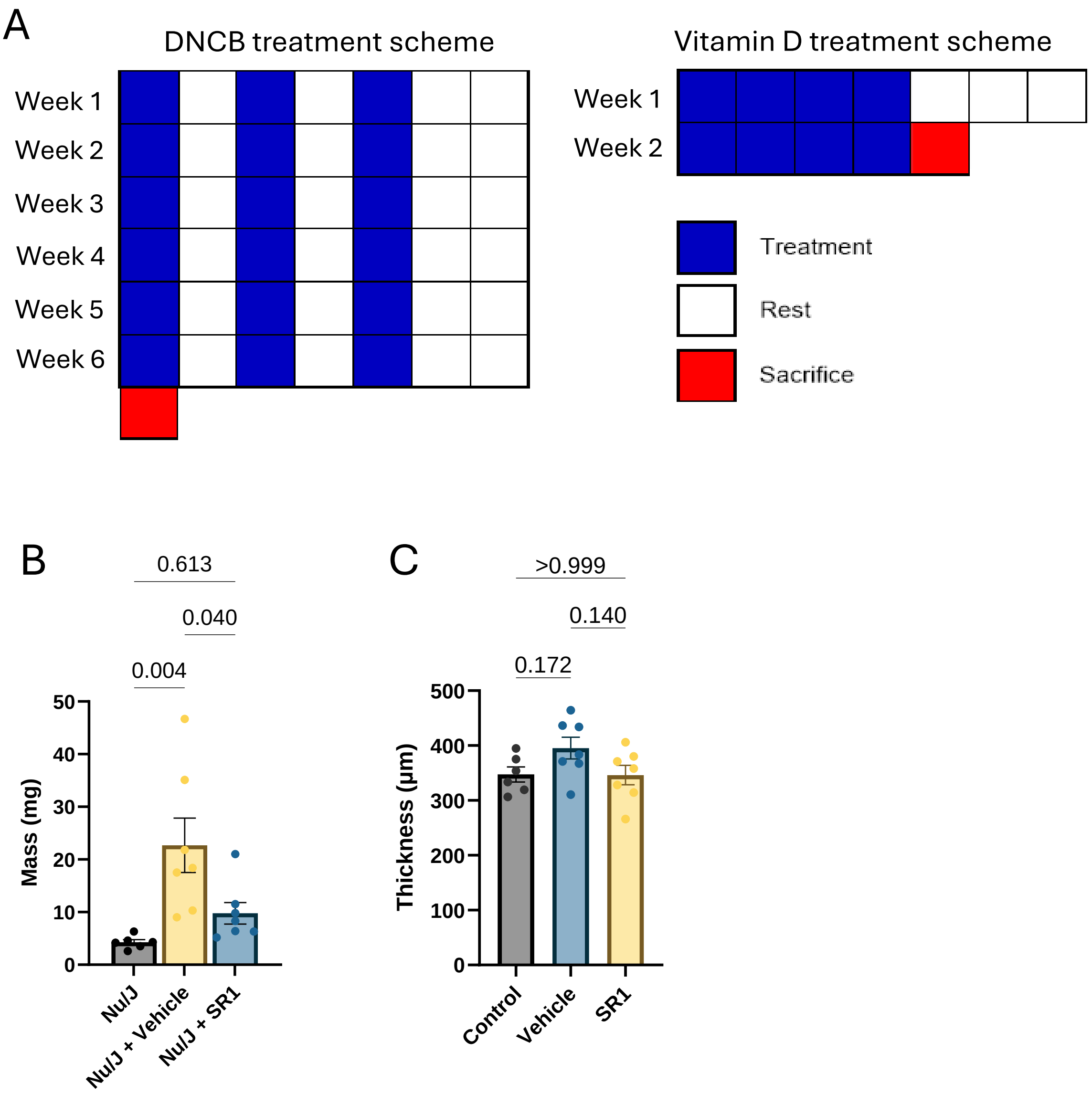

A
DNCB treatment scheme
Week 1
Week 2
Week 3
Week 4
Week 5
Week 6
Vitamin D treatment scheme
Week 1
Week 2
Treatment
Rest
Sacrifice
B
0.613
0.040
0.004
Mass (mg)
50
40
30
20
10
0
Nu/J
Nu/J + Vehicle
Nu/J + SR1
C
>0.999
0.140
0.172
Thickness (µm)
500
400
300
200
100
0
Control
Vehicle
SR1
